# CRY-NLRP3 complexes define a circadian checkpoint controlling inflammasome activation

**DOI:** 10.64898/2026.02.26.708242

**Authors:** Léa Bardoulet, Delphine Burlet, Hubert Leloup, François Virard, Sabine Hacot, Elsa G. Guillot, Nouhaila El Kasmi, Camille Cosson, Lyvia Moudombi, Marie-Cécile Michallet, Arnaud Bonnafoux, Mélina Gautier, Agnès Tissier, Katja A. Lamia, Bénédicte F Py, Virginie Petrilli, Anne-Laure Huber

## Abstract

Innate immune sensors such as the NLRP3 inflammasome can trigger inflammatory responses within minutes, raising the question of how circadian clocks influence such rapid decisions. Here, we identify a protein-level circadian checkpoint that links core clock components to NLRP3 inflammasome activation. We show that NLRP3 associates with the circadian repressors CRY1 and CRY2, forming oscillatory complexes that restrain inflammasome activation and rapidly dissociate upon stimulation. Pharmacological stabilization of CRY proteins preserves CRY-NLRP3 association and attenuates inflammasome assembly, IL-1β secretion and pyroptotic cell death in primary human macrophages. In synchronized macrophages, both NLRP3 inflammasome activation and its inhibition by the NLRP3 inhibitor MCC950 vary with circadian time. Finally, a subset of NLRP3 variants reported in cohorts of patients with Cryopyrin-Associated Periodic Syndromes (CAPS), a group of hereditary fever syndromes caused by mutations in NLRP3, weaken CRY binding and are associated with altered time-of-day patterns of inflammasome activation and MCC950 responsiveness. Together, these findings define CRY-NLRP3 complexes as a circadian checkpoint that modulates inflammasome activity and drug response, revealing time of day as a critical dimension of NLRP3-driven inflammation.

## Introduction

Circadian clocks enable organisms to anticipate daily environmental changes and adjust physiological processes accordingly. In mammals, circadian regulation extends deeply into immune function, shaping leukocyte trafficking, cytokine production, and pathogen sensing (1–4). In macrophages, intrinsic clocks drive daily oscillations in gene expression, inflammatory mediator release, and phagocytic activity (5–8). At the molecular level, circadian rhythms are generated by a transcription-translation feedback loop in which the basic helix-loop-helix ARNT like 1(BMAL1) together with circadian locomoter output cycles protein kaput (CLOCK) drives Period (Per1-3) and Cryptochrome (Cry1-2) expression, and PER/CRY proteins feedback to inhibit BMAL1-CLOCK activity, generating 24-hour oscillations (9,10). Auxiliary loops involving REV-ERB and ROR stabilize this core oscillator and extend circadian control to many downstream genes, including immune regulators (11–13).

Accumulating evidence indicates that core clock components actively restrain inflammatory signaling. Cry1/2-deficient mice display elevated cytokine production and increased susceptibility to inflammatory disease, while macrophages lacking Bmal1 exhibit increased IL-1β production and mitochondrial stress (7,14–16). CRY proteins are central repressors of circadian transcription and are tightly regulated by phosphorylation- and ubiquitin-dependent turnover (17,18). Beyond their canonical role in the clock, CRYs also function as scaffolds that recruit E3 ubiquitin ligases to regulate the stability of interacting proteins independently of gene expression (19). These properties position CRY proteins as potential post-translational regulators of signaling complexes. Together, these observations raise the possibility that clock proteins engage innate immune pathways through protein-level mechanisms.

To determine whether circadian clock components physically associate with innate immune sensors, we assessed interactions between core clock proteins and selected pattern recognition receptors (PRRs). This analysis revealed a previously unrecognized association between CRY proteins and NLR family pyrin domain containing 3 (NLRP3).

NLRP3 is a cytosolic pattern recognition receptor of the NOD-like receptor family that functions as the sensor component of the NLRP3 inflammasome, a multiprotein innate immune complex. As a sensor of cellular stress, NLRP3 responds to pathogen- and damage-associated signals by oligomerizing and assembling with the adaptor protein ASC, which recruits caspase-1 to form an active inflammasome complex (20–23). Inflammasome assembly results in caspase-1 activation, maturation of the pro-inflammatory cytokines IL-1β and IL-18, and cleavage of gasdermin-D (GSDMD), whose N-terminal fragment forms membrane pores that enable cytokine release and trigger pyroptotic cell death (24,25).

Excessive inflammasome activation is highly pathogenic, dysregulated NLRP3 signaling underlies monogenic autoinflammatory disorders such as Cryopyrin-Associated Periodic Syndromes (CAPS) and has also been implicated in prevalent chronic inflammatory diseases including type 2 diabetes, atherosclerosis, and Alzheimer’s disease (26). Accordingly, NLRP3 activity is tightly regulated at multiple levels, from transcriptional priming that controls NLRP3 expression to post-translational mechanisms, including ubiquitination and phosphorylation, that modulate NLRP3 stability and its capacity to assemble into a functional inflammasome (27–31). Circadian regulation has been linked to NLRP3-driven inflammation primarily through transcriptional and metabolic programs. Circadian factors such as REV-ERBα and BMAL1 influence inflammasome outputs by shaping inflammatory gene expression and cellular metabolism, including mitochondrial function and redox balance (32–35).

While these transcriptional and metabolic mechanisms clearly contribute to circadian regulation of inflammation, they do not establish whether circadian timing is encoded directly within the inflammasome complex itself. The identification of a CRY–NLRP3 association raises the possibility that clock components may modulate inflammasome assembly or activation through protein-level mechanisms. Determining the functional consequences of this interaction is therefore critical to understand how circadian timing intersects with innate immune signaling. Here, we identify a post-translational mechanism by which CRY1 and CRY2 interact with NLRP3 to restrain inflammasome activation. We demonstrated that CRY-NLRP3 complexes oscillate over circadian time and undergo rapid dissociation upon inflammasome stimulation. Pharmacological stabilization of CRY proteins preserves CRY-NLRP3 association and attenuates inflammasome assembly and pyroptotic cell death in primary human macrophages. Consistent with this protein-level checkpoint, both NLRP3 inflammasome activity and the efficacy of the NLRP3 inhibitor MCC950 vary according to circadian timing. Moreover, NLRP3 variants identified in CAPS-associated contexts weaken CRY binding and alter CRY stability, thereby modifying how inflammasome activation and its pharmacological inhibition change over circadian time. Together, these findings establish CRY–NLRP3 complexes as a dynamic circadian checkpoint that links clock function to inflammasome activation and drug responsiveness.

## Results

### NLRP3 inflammasome activity is anti-phasic to CRY levels in primary human macrophages

Circadian regulation of NLRP3 inflammasome activity has been reported (33–35), yet the molecular mechanisms that gate rapid inflammasome activation in a time-of-day-dependent manner at the protein level remain unclear. We isolated CD14⁺ monocytes from five healthy donors and differentiated them into macrophages (Figure S1A). Cells were synchronized using a time-shifted serum-shock protocol in which serum-shock was applied every 4 h to enable multiple times after synchronization to be assayed in parallel on the same plate (Figure S1B). Unless otherwise indicated, samples were collected between 12 and 48 h after synchronization. Following LPS priming and nigericin stimulation, a microbial toxin that triggers K⁺ efflux and activates NLRP3, inflammasome outputs displayed robust time-of-day dependence in primary human macrophages. Pronounced differences across times after synchronization in ASC speck formation, IL-1β release, and inflammasome-associated cell death (pyroptosis), with peak activity 17-18 hr and a trough around 29-30 hr post-synchronisation (Figure 1A-C). Caspase-1 activity showed only modest variation across circadian time but followed the same overall trend as the other inflammasome readouts (Figure 1D). We monitored NLRP3, CRY1, and CRY2 protein abundance across this window and refer to phases with high or low CRY expression as CRY-high or CRY-low, respectively (Figure 1E). Interestingly, we noticed that the CRY-high phase correlates with minimal inflammasome activity, consistent with an inverse relationship between CRY abundance and NLRP3 inflammasome responsiveness. This anti-phasic pattern between the CRY-high state and inflammasome output suggests that CRY proteins might functionally interact with NLRP3 at the protein level, rather than acting solely through transcriptional regulation.

**Figure 1.**
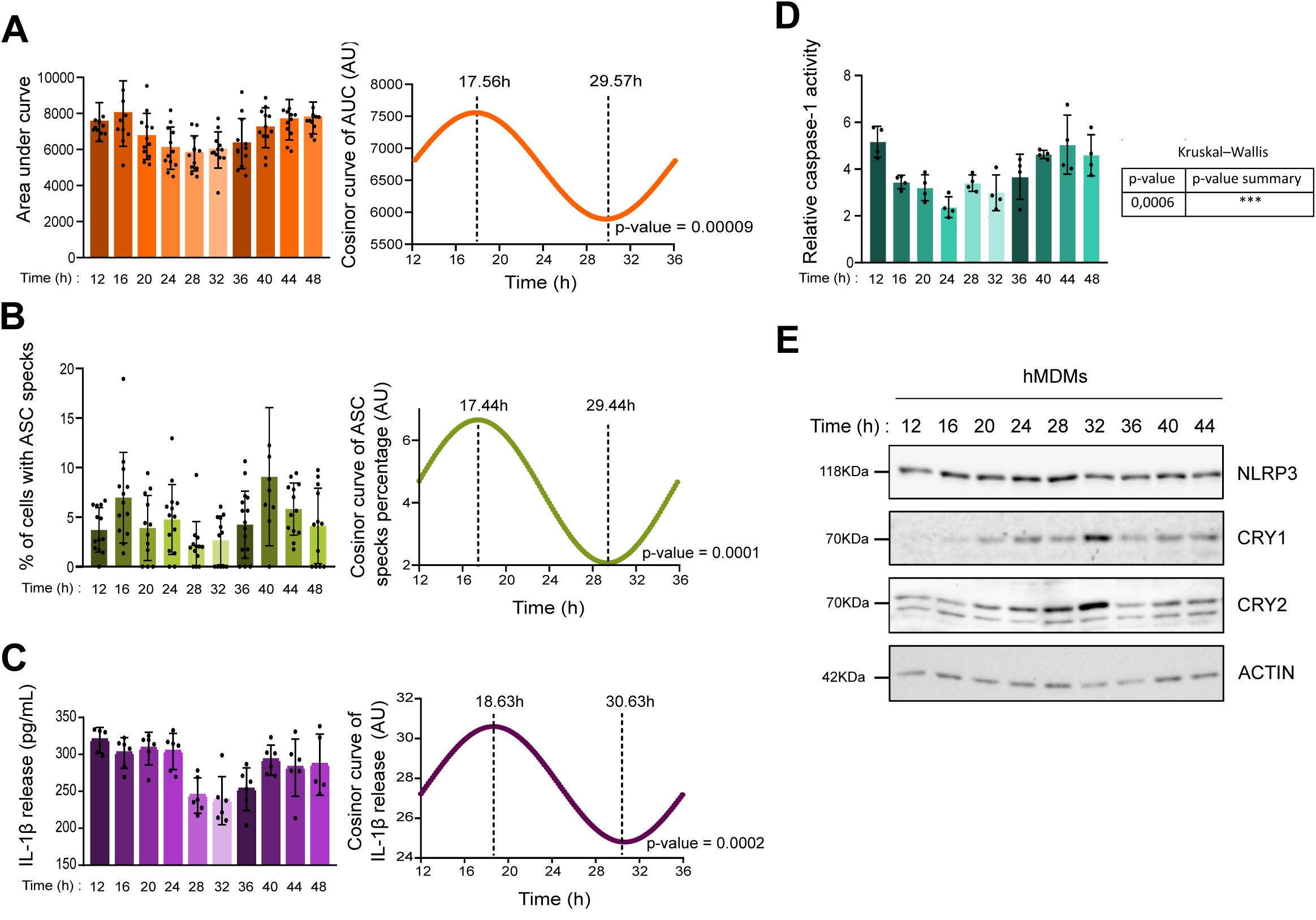
NLRP3 inflammasome activity is anti-phasic to CRY levels in primary human macrophages. (A - E) hMDMs were synchronized by a 2-h serum shock. Cells were primed with LPS (0.5 µg/mL, 4 h) and activated with nigericin (20 µM, 30-240 min) at different time post synchronization (from CT12 to CT48). (A) Cell death monitored by DRAQ7 incorporation using an Opera Phenix HCS microscope. Area under the curve (AUC) values were quantified across time after synchronization (pooled data from three independent experiments), shown as mean ± SD. (B) ASC-speck-positive cells quantified at each circadian time point from three independent experiments, each performed with four experimental replicates (12 wells per condition); with ≥200 cells analyzed per condition, mean ± SD. (C) IL-1β secretion measured by ELISA in supernatants from hMDMs treated with nigericin (20 µM, 240 min) at each time point after synchronization. Data are from three independent experiments, each performed with two experimental replicates (6 wells per condition), shown as mean ± SD. (A, B, C) Right panel: Cosinor fits of raw data generated with COSINOR online software, showing acrophase and bathyphase; p values calculated by Cosinor analysis. (D) Caspase-1 activity normalized to Crystal Violet (CV) staining. Data represent four independent experiments, each performed with six technical replicates, shown as mean ± SD. Kruskal-Wallis test, p = 0.0006 (***). (E) hMDMs were collected every 4 h from 12 h to 48 h after synchronization, following LPS priming (0.5 µg/mL, 4 h). Protein expression of NLRP3, CRY1 and CRY2 were analyzed by immunoblotting; actin served as a loading control.

### CRY proteins form complexes with NLRP3 and display time-of-day-dependent association in primary human macrophages

Beyond their canonical role in circadian transcription, CRYs have been reported to assemble with additional partners in protein homeostasis pathways (19). These observations, together with the anti-phasic relationship between CRY abundance and inflammasome outputs, raised the possibility that core clock components might influence the NLRP3 inflammasome at the post-translational level. We therefore examined whether NLRP3 interacts with CRY proteins in primary human macrophages. We first tested this interaction in THP-1 macrophages under endogenous conditions by co-immunoprecipitation, comparing cells synchronized by serum shock 6hr post synchronization, when CRY expression peaks, with unsynchronized cells. In both conditions, CRY2 was recovered with endogenous NLRP3, while no interaction was detected in NLRP3 knockout controls (Figure 2A). We further validated these interactions in transfected HEK293T cells, where FLAG-NLRP3 was co-expressed with CRY1 (Figure S2A) or CRY2 (Figure S2B) and co-immunoprecipitated with FLAG-NLRP3. Proximity ligation assays (PLA) further confirmed the association between NLRP3 and CRY proteins in multiple cellular models. In THP-1 macrophages, we observed robust PLA signals for both CRY1-NLRP3 and CRY2-NLRP3, whereas no signal was observed in NLRP3 knockout cells, confirming assay specificity (Figure 2B; Figure S2C, D). Similar results were obtained in U937 monocytes, where CRY1-NLRP3 and CRY2-NLRP3 complexes were detected only in wild-type cells (Figure S2E-G). In hMDMs, we confirmed that NLRP3 associates with CRY1 and CRY2 in unsynchronized hMDMs (Figure 2C, S2I-J). Control reactions with single antibodies produced no signal in any of the systems tested, excluding non-specific amplification (Figure S2H-I). Overall, the data indicate that CRY1 and CRY2 interact with NLRP3 in myeloid cells. We finally asked whether CRY proteins interact with other inflammasome components. In HEK293T cells, neither CRY1 nor CRY2 co-immunoprecipitated with the inflammasome adaptor ASC (Figure S2K-L). Taken together, these results indicate that CRY1 and CRY2 selectively form complexes with NLRP3 in different cellular models.

**Figure 2.**
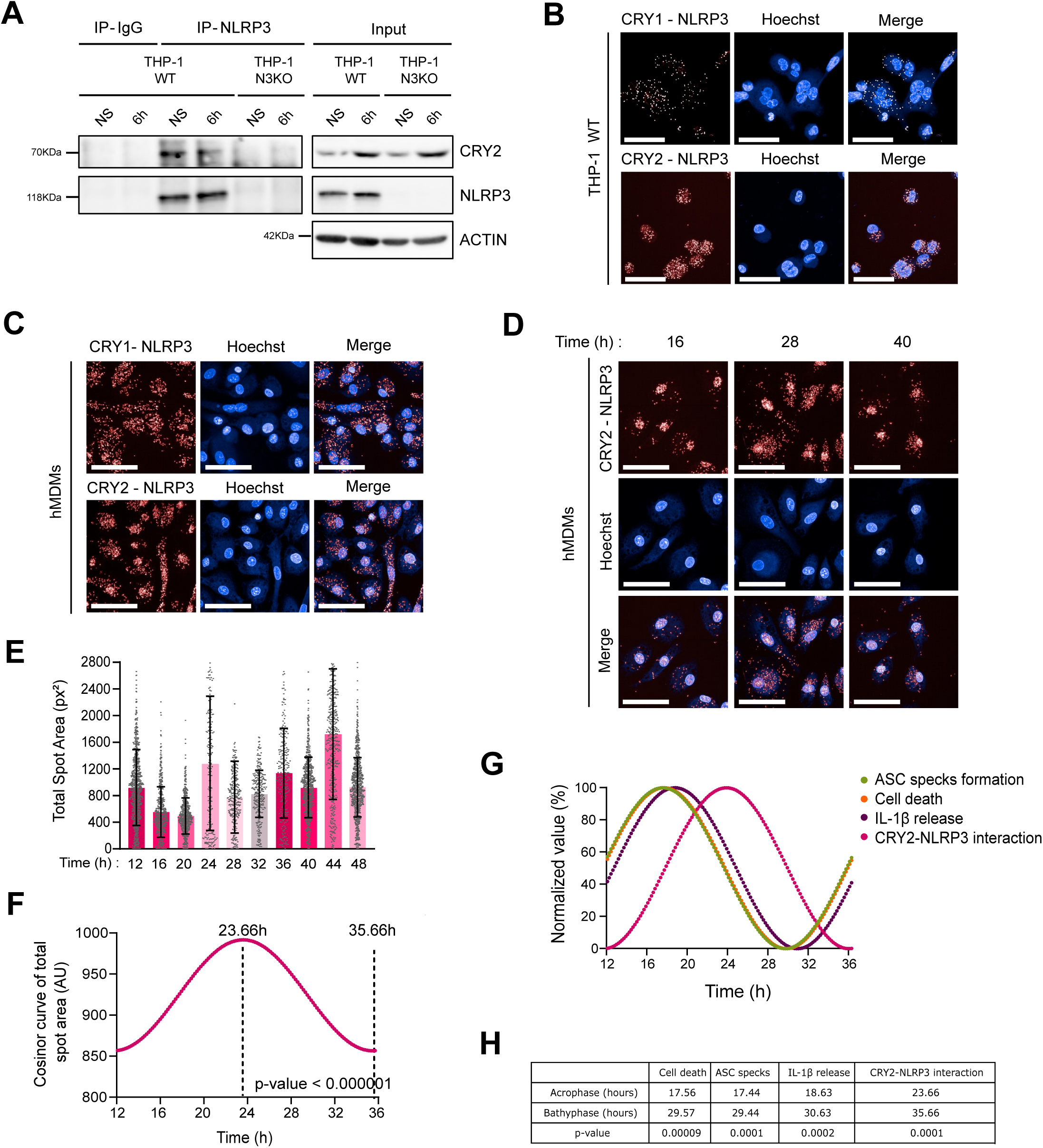
CRY proteins form complexes with NLRP3 and display time-of-day-dependent association in primary human macrophages. (A) Endogenous NLRP3 was immunoprecipitated from THP-1 WT or NLRP3 knockout (N3KO) cells using anti-NLRP3 antibodies. Immunoprecipitation was performed 6 h after serum shock synchronization or in unsynchronized (NS) conditions, as indicated. IgG served as a control. CRY2 co-immunoprecipitation was assessed by immunoblotting; actin was used as a loading control (n=2). (B-C) Representative proximity ligation assay (PLA) images showing CRY1- NLRP3 or CRY2-NLRP3 interactions (red) in untreated THP-1 WT cells (n=4) (B) and hMDMs (n>5) (C). Nuclei were stained with Hoechst; images were acquired on an Opera Phenix HCS microscope using a 60× objective; scale bar, 50 μm. (D) Representative PLA images (red) showing CRY2- NLRP3 interactions in hMDMs at 16 h, 28 h, and 40 h after synchronization and following LPS priming. Nuclei were stained with Hoechst. Images were acquired on an Opera Phenix HCS microscope with a 40× water objective; scale bar, 50 µm; n > 1,000 cells/condition. (E) Quantification of PLA total spot area per cell from (D). (F) Cosinor analysis of data in (E) showing circadian oscillation with acrophase and bathyphase indicated. (G) Cosinor curves for cell death, ASC speck formation, IL-1β secretion, and CRY2- NLRP3 interactions plotted together. (H) Summary table reporting acrophase, bathyphase, and cosinor p values for panels (Fig 1A, B, and Fig 2F).

We next examined whether CRY-NLRP3 interactions oscillate across circadian time. PLA revealed marked temporal variation in CRY2-NLRP3 complexes, as illustrated by representative confocal images (Figure 2D). Quantification confirmed significant circadian changes in CRY2-NLRP3 association across circadian time (Figure 2E). Cosinor analysis demonstrated robust rhythmicity, with peak interaction between 23-24 hr post-synchronization (Figure 2F).

To evaluate donor-to-donor robustness, we quantified CRY2-NLRP3 PLA signal at each time after synchronization for each donor individually (Figure S3A-K). Because PLA values were not normally distributed and donor-to-donor differences in amplitude and phase precluded pooling, each donor was analyzed using Kruskal-Wallis analysis. This analysis confirmed significant time-dependent variation in CRY2-NLRP3 PLA signal within each donor (Figure S3K). Rhythmicity was supported by cosinor analysis (Figure 2F, S3B,D,F,H,J). Importantly, applying the same PLA acquisition and analysis pipeline to an unrelated protein pair (ATM-ATM) in the same donor revealed no time-dependent variation (Figure S3L), supporting the specificity of CRY2-NLRP3 oscillations.

To correlate CRY2-NLRP3 interactions with inflammasome activity over circadian time, we superposed cosinor curves for cell death, ASC speck formation, IL-1β secretion, and CRY2-NLRP3 proximity (Figure 2G,H). This integrated representation revealed an inverse relationship between CRY2-NLRP3 complex formation and inflammasome activity, with maximal cell death, ASC speck formation, and IL-1β secretion occurring when CRY2-NLRP3 association was low. Together, these results identify CRY2-NLRP3 complexes as a circadian, post-translational checkpoint that restrains inflammasome activation. By oscillating in antiphase with inflammasome output, CRY-NLRP3 interactions provide a mechanism by which circadian phase sets time-dependent thresholds for NLRP3 responsiveness.

### Inflammasome activation disrupts circadian CRY-NLRP3 complexes in primary human macrophages

Because inflammasome outputs peak at CRY-low phases and CRY2-NLRP3 complex abundance is anti-phasic to inflammasome activity, we asked whether acute inflammasome activation alters CRY protein levels and CRY-NLRP3 complex abundance. We first assessed CRY protein levels in hMDMs and THP-1 cells following LPS priming and acute nigericin stimulation and observed reduced CRY1 and CRY2 abundance (Figure 3A, S4A). We then tested whether this decrease was accompanied by reduced CRY-NLRP3 proximity. In both cell lines, PLA revealed a marked reduction in CRY1-NLRP3 and CRY2-NLRP3 signal after nigericin stimulation, as shown by representative confocal images and quantification (Figure 3B-D; S4B-D). This reduction is consistent with decreased CRY-NLRP3 complex abundance, which may reflect reduced CRY protein levels and/or weakened association between CRY and NLRP3. Complementary co-immunoprecipitation assays in HEK293T cells further supported a nigericin-dependent reduction in CRY-NLRP3 association (Figure S4E,F).

**Figure 3.**
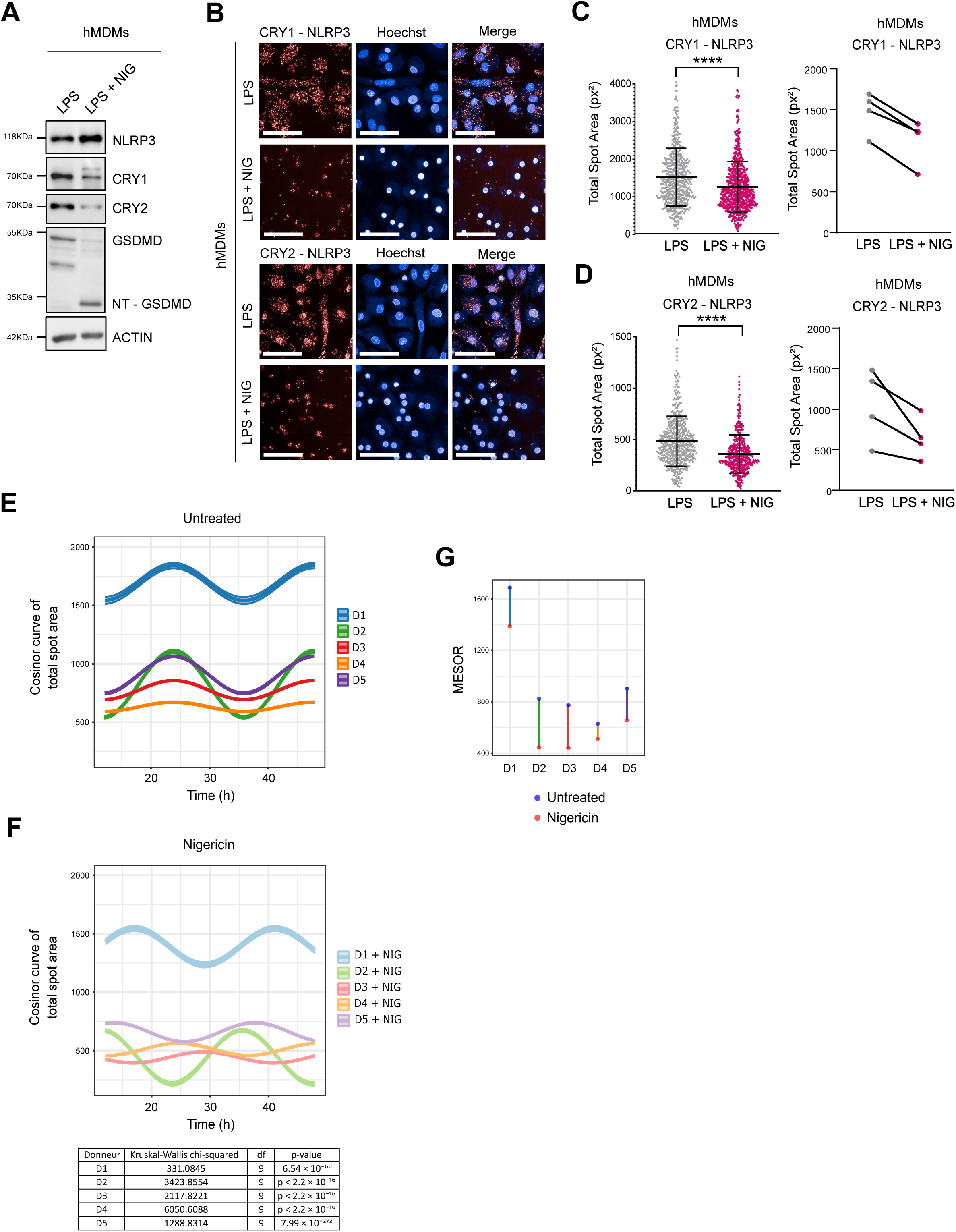
Inflammasome activation disrupts circadian CRY-NLRP3 complexes in primary human macrophages. (A) Immunoblot analysis of NLRP3, CRY1, CRY2, and GSDMD in hMDMs (n=6) treated with LPS (0.5 µg/mL, 4 h) with or without nigericin (10 µM, 1 h). Actin served as a loading control. (B) Representative PLA images (red) showing CRY1-NLRP3 or CRY2-NLRP3 interactions in hMDMs (N=6) primed with LPS (0.5 µg/mL, 4 h) and left untreated or activated with nigericin (20 µM, 1 h). Nuclei were stained with Hoechst; images were acquired on an Opera Phenix HCS microscope using a 40× objective; scale bar, 50 µm. (C-D) Quantification of PLA total spot area per cell for CRY1-NLRP3 (C) and CRY2-NLRP3 (D). ≥500 cells per condition; mean ± SEM; Mann-Whitney test, ****p < 0.0001. (E and F) Pooled Cosinor fits of CRY2-NLRP3 PLA signal over time from the five donors shown in Figure S5, either untreated (E) or treated with nigericin (F). (G) Lollipop plots showing MESOR values of CRY2-NLRP3 interactions corresponding to the cosinor fits in (E-F). Blue indicates untreated cells; pink indicates nigericin-treated cells.

Given that CRY2-NLRP3 complexes oscillate over time (Figure 2), and considering that nigericin triggers complex dissociation, we investigated the impact of this treatment on CRY2-NLRP3 dynamics. To compare donors with phase-shifted rhythms, we aligned untreated traces per donor and applied the same offsets to the corresponding nigericin traces. Nigericin consistently reduced the overall CRY2-NLRP3 interaction signal across donors and generally dampened the oscillatory pattern (Figure 3E-F; Figure S5).

To quantify these effects, we extracted three parameters for each donor: mean level (MESOR), amplitude, and phase. Nigericin lowered the mean CRY2-NLRP3 interaction level in all donors (MESOR; Figure 3G), showing that inflammasome activation consistently reduces CRY2-NLRP3 complex abundance. Amplitude and phase responses were more heterogeneous (Figure S5K-L): in some donors the rhythm was strongly dampened, whereas in others the timing or strength of the peak changed without a clear loss of rhythmicity. Importantly, however, every donor showed a decrease in the average level of CRY2-NLRP3 interaction, indicating a shared core response to inflammasome activation, with donor-specific differences in how the circadian pattern is modulated.

### Pharmacological stabilization of CRY attenuates NLRP3 inflammasome activation

Because acute inflammasome activation reduces CRY abundance and destabilizes CRY-NLRP3 complexes, we asked whether pharmacological stabilization of CRY proteins could preserve CRY-NLRP3 complexes and thereby restrain NLRP3 inflammasome activation. We tested KL001 and KL044, previously described as stabilizers that preferentially target either CRY1 or CRY2 (36,37), depending on the cellular context, and TH301, which preferentially stabilizes CRY2 (38). Immunoblotting in hMDMs confirmed that KL001 and KL044 increased CRY2 abundance with minimal effects on CRY1, while TH301 robustly stabilized CRY2 as expected (Figure S6A-C).

We next examined the functional consequences of CRY stabilization on nigericin-induced cell death. Live-cell imaging of DRAQ7 incorporation showed that KL001 pretreatment consistently slowed the initiation and reduced the extent of cell death, as reflected by the area under the curve across multiple donors (Figure 4A). KL044 and TH301 produced similar effects, though less pronounced (Figure 4B; Figure S6D). Despite inter-donor variability, all three compounds showed the same overall trend.

**Figure 4.**
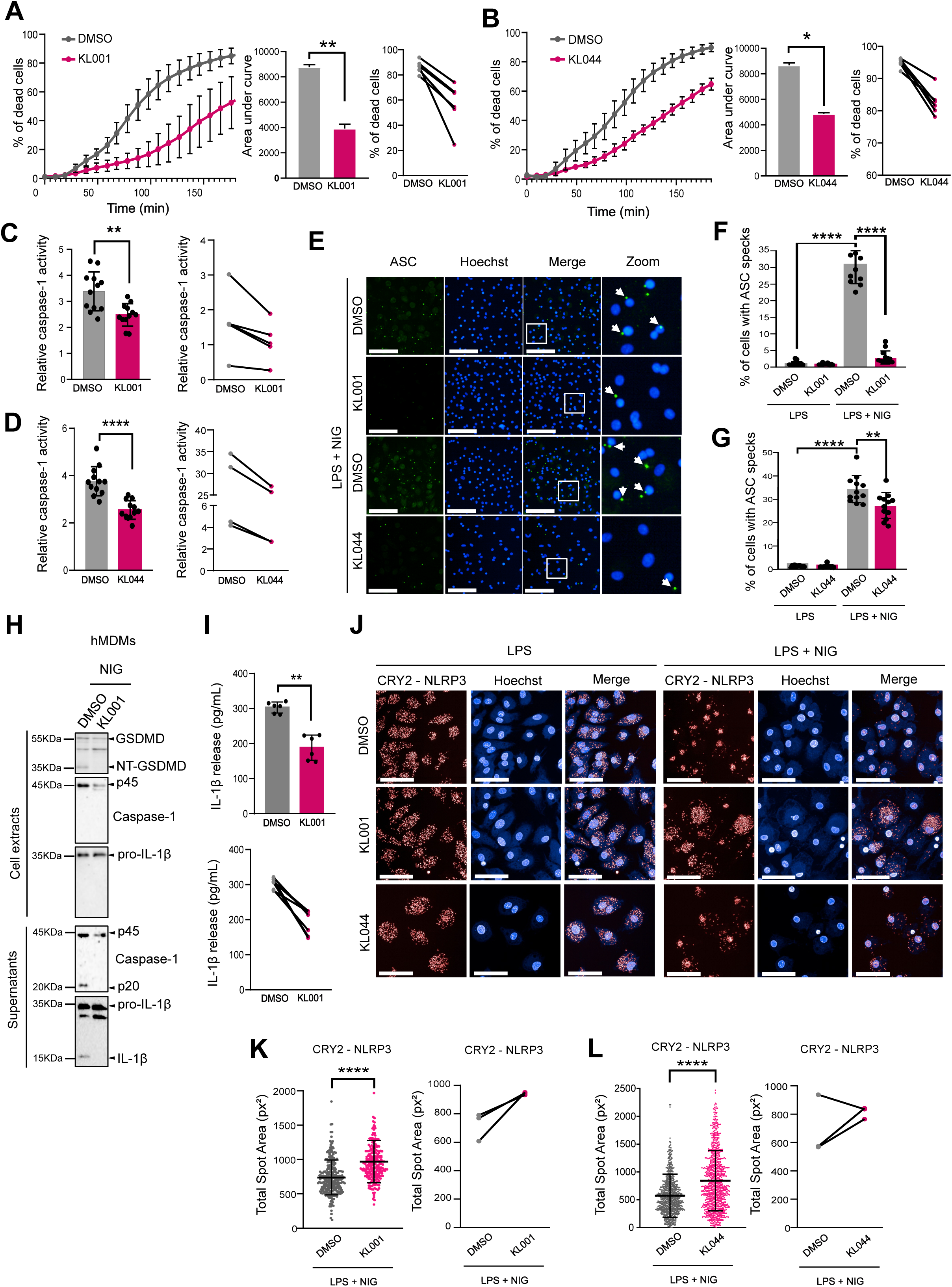
Pharmacological stabilization of CRY proteins attenuates NLRP3 inflammasome activation in primary human macrophages. (A-G) hMDMs were differentiated with M-CSF (7 d) and pretreated for 48 h with DMSO or CRY stabilizers KL001 (5 µM) (A,C,E,F) or KL044 (5 µM) (B,D,E,G), then primed with LPS (0.5 µg/mL, 4 h) and activated with nigericin (20 µM). (A,B) Time course of cell death measured by DRAQ7 incorporation (Opera Phenix). Left: kinetic curves of DRAQ7 signal over time. Middle: quantification of cell death as area under the curve (AUC) (GraphPad Prism), shown as mean ± SD. Upper: final point from paired individual values from six independent experiments, each represented by three pooled technical replicates (18 total points). Statistical comparisons were performed using a paired Mann-Whitney test (*p < 0.03, **p < 0.005). (C, D) Caspase-1 activity normalized to CV staining. Left: pooled quantification shown as mean ± SD from six independent experiments, each performed with two technical replicates (12 total values). Lower: paired individual values from the six independent experiments. Statistical comparisons were performed using a paired Mann-Whitney test (**p < 0.005, ****p < 0.0001). (E) Representative confocal images of ASC specks (green) in hMDMs preteated with DMSO, KL001 (5 µM) or KL044 (5 µM), primed with LPS (0.5 µg/mL, 4 h) and left untreated or activated with nigericin (20 µM, 1.5 h). Nuclei were stained with Hoechst; images were acquired on an Opera Phenix HCS microscope using a 20× air objective; scale bar, 50 µm; arrows indicate ASC specks. (F, G) Quantification of ASC-speck-positive shown as mean ± SD from three independent experiments, each performed with three experimental replicates (9 wells per condition) with ≥200 cells analyzed per condition. Statistical comparisons were performed using a paired Mann-Whitney test (**p = 0.0068, ****p < 0.0001). (H) hMDMs were pre-treated with DMSO or KL001 (5 µM), primed with LPS (0.5 µg/mL, 4 h) and treated with nigericin (20 µM) for 3 h. Levels of GSDMD, caspase-1, and IL-1β were analyzed by immunoblotting in cell lysates and corresponding culture supernatants (n = 2). (I) IL-1β secretion measured by ELISA in supernatants from hMDMs pretreated with KL001 (5 µM), primed with LPS (0.5 µg/mL, 4 h) and treated with nigericin (20 µM, 3 h). Top: pooled quantification from six measurements obtained across four independent experiments, shown as mean ± SD. Bottom: paired individual values from the four independent experiments. Statistical comparisons were performed using an unpaired Mann-Whitney test. (J) Representative PLA images (red) showing CRY1- NLRP3 or CRY2-NLRP3 interactions in hMDMs pretreated with DMSO, KL001 (5 µM), or KL044 (5 µM), primed with LPS (0.5 µg/mL, 4 h) and left untreated or activated with nigericin (20 µM, 1 h). Nuclei were stained with Hoechst; images were acquired on an Opera Phenix HCS microscope using a 40× water objective; scale bar, 50 µm. (K-L) Quantification of PLA total spot area per cell for CRY2-NLRP3 after KL001 treatment (K) or KL044 treatment (L), as shown in (J). Left: representative experiment showing three wells per condition with ≥1,000 cells analyzed per well. Right: paired summary of three independent experiments. Data are shown as paired dots with mean ± SD. Statistical significance was determined using a paired Mann-Whitney test (****p < 0.0001).

We next assessed inflammasome activity using caspase-1 activation, ASC speck formation, and interleukin-1β (IL-1β) processing as readouts. Caspase-1 activity was significantly reduced by KL001, KL044, and TH301 (Figure 4C,D; Figure S6E). In addition, ASC speck formation showed marked reduction upon KL001 treatment and a milder reduction upon KL044 and TH301 (Figure 4E-G; Figure S6F-H), consistent with reduced inflammasome assembly in the presence of CRY stabilizers. Accordingly, KL001 (Figure 4H,I) and TH301 (Figure S6I,J) decreased caspase-1 auto-processing and IL-1β cleavage and secretion following nigericin stimulation.

Since CRY stabilization reduced nigericin-induced inflammasome outputs, we next asked whether this protection was associated with sustained CRY-NLRP3 interactions. We therefore performed PLA in primary hMDMs pretreated with CRY stabilizers (Figure 4J-L; Figure S6K-M). After nigericin stimulation, KL001 and KL044-treated cells retained more CRY-NLRP3 complexes than DMSO controls. Across conditions, preservation of CRY-NLRP3 complexes coincided with reduced cell death, supporting the idea that CRY binding provides an additional layer of control over NLRP3 activation in primary human macrophages.

To confirm that the observed effects were largely NLRP3-dependent, we repeated key assays in the presence of the selective NLRP3 inhibitor MCC950 (39). MCC950 pretreatment consistently reduced nigericin-induced cell death (Figure S6N,O) and markedly decreased caspase-1 activation (Figure S6P), confirming that the measured outputs predominantly reflect NLRP3 inflammasome activity.

### Nigericin promotes FBXL3-dependent ubiquitination of CRY proteins

Since stabilizing CRY proteins attenuated inflammasome activation, we asked how nigericin drives the loss of CRY proteins. To test whether the decrease in CRY abundance involves ubiquitin-mediated destabilization, HEK293T cells were transfected with CRY1 (Figure 5A) or CRY2 (Figure 5B) together with HA-tagged ubiquitin, with or without NLRP3. After nigericin stimulation, ubiquitinated proteins were isolated by HA pulldown, and CRY was detected by immunoblot. Nigericin markedly increased the amount of CRY recovered in the HA-ubiquitin IP, whereas co-expression of NLRP3 reduced this signal, suggesting reduced availability of CRY for ubiquitination when associated with NLRP3. Given the established role of the SCF E3 ligase receptor FBXL3 in regulating CRY stability (17,18), we hypothesized that nigericin-induced ubiquitination could result from enhanced FBXL3 binding. Consistent with this idea, co-immunoprecipitation experiments showed a clear increase in FBXL3-CRY association following nigericin stimulation (Figure 5C,D). To validate the role of FBXL3 in promoting CRY ubiquitination, we compared wild-type FBXL3 with a ΔF-box mutant that cannot assemble into an active SCF complex (19). In our assay, only wild-type FBXL3 promoted nigericin-induced ubiquitination, whereas the ΔF-box mutant failed to do so (Figure 5E,F). To further test the requirement for FBXL3 engagement, we analyzed CRY2 mutants that specifically disrupt FBXL3 binding (F428D, I499D, L517D) (40). These mutants were expressed in HEK293T cells together with FBXL3 and subjected to nigericin treatment over a time course. Wild-type CRY2 levels decreased by ∼50% within one hour of stimulation, whereas all three mutants remained markedly more stable, demonstrating that nigericin-stimulated CRY2 destabilization requires intact FBXL3 binding (Figure 5G).

**Figure 5.**
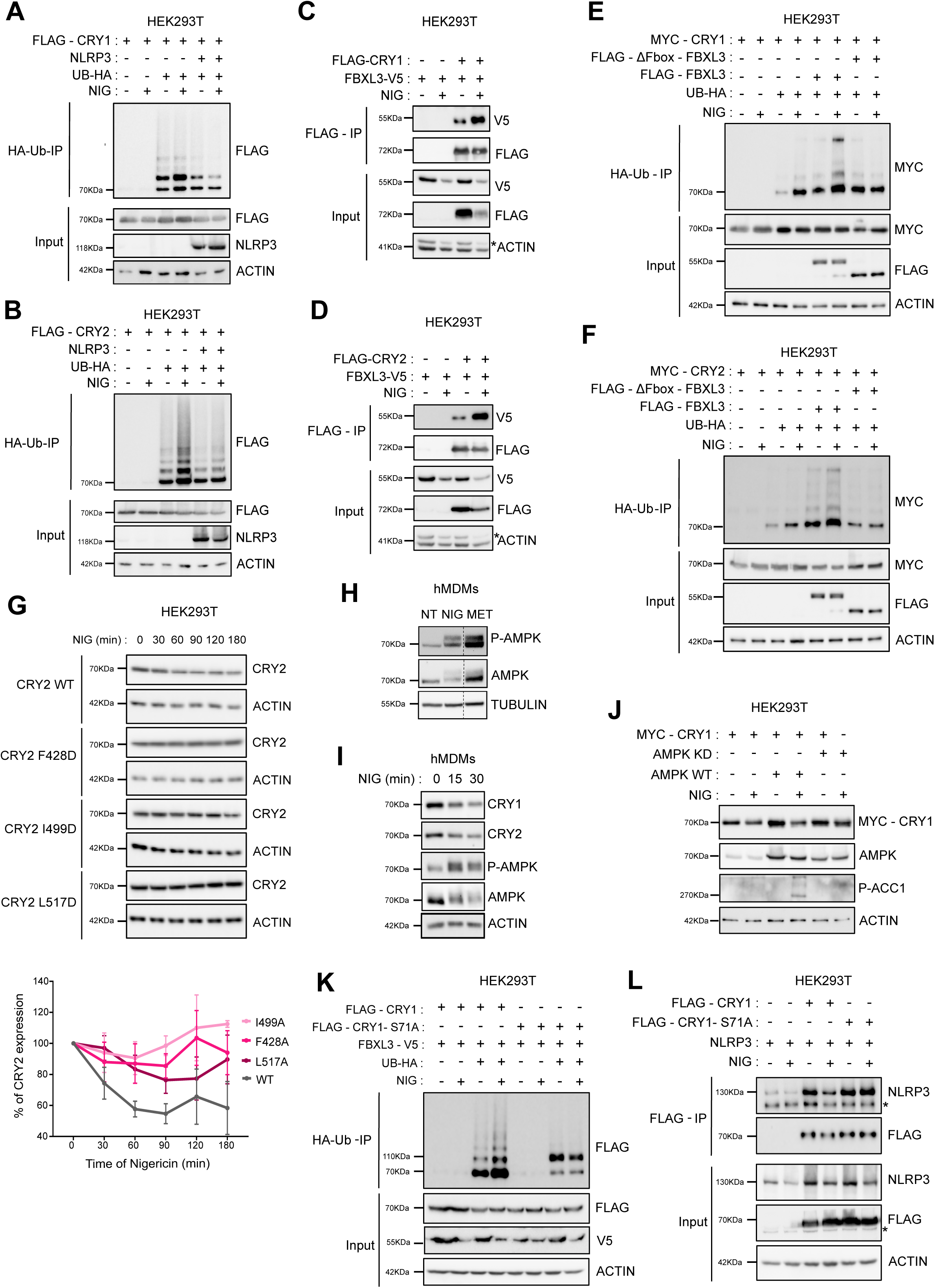
Nigericin promotes FBXL3 recruitment to CRYs, enhancing ubiquitination and degradation. (A-B) HEK293T cells were transfected with the indicated constructs Ub-HA, NLRP3, FLAG-CRY1 or FLAG-CRY2 (N=4); or with (C-D) FBXL3-V5, FLAG-CRY1 or FLAG-CRY2 (n=3). Cells were treated with nigericin (10 µM, 1 h) before HA-Ub or anti-FLAG immunoprecipitation respectively. (E-F) Ub-HA, MYC-CRY1 (E) or MYC-CRY2 (F) were transfected in HEK293T cells with FBXL3-V5, FLAG-FBXL3, or FLAG-ΔF-box-FBXL3. Cells were treated with nigericin (10 µM, 1 h) before HA-Ub immunoprecipitation. CRY1 and CRY2 ubiquitination was assessed by immunoblotting (n=2). (G) HEK293T cells expressing WT or FBXL3-binding-deficient CRY2 mutants (F428D, I499D, L517D) were treated with nigericin (10 µM, 0-180 min). CRY2 expression was analyzed by immunoblotting and quantified (lower panel) using the Image Lab software (Bio-Rad) (n=3). (H) HEK293T cells were treated with nigericin (10 µM, 1 h), or metformin (20 mM, 1 h). Phosphorylated AMPK (P-AMPK) and total AMPK levels were assessed by immunoblotting. Tubulin served as a loading control (n=3). (I) hMDMs were treated with nigericin (20 µM) for 0, 15, or 30 min. CRY1, CRY2, P-AMPK, and AMPK levels were analyzed by immunoblotting. Actin served as a loading control (n=3). (J) Whole-cell lysates from HEK293T cells transfected with AMPK WT, AMPK kinase-dead (KD), and MYC-CRY1 were analyzed by immunoblotting for MYC-CRY1, AMPK and P-ACC1. Actin served as a loading control. (n=2) (K) HEK293T cells expressing WT or S71A CRY1 with V5-FBXL3 and HA-Ub were subjected to HA-Ub immunoprecipitation after nigericin treatment (10 µM, 1 h); CRY1 ubiquitination was analyzed by immunoblotting (n=3). (L) HEK293T cells expressing NLRP3 and WT or S71A CRY1 were treated with nigericin (10 µM, 1 h). Anti-FLAG IP and immunoblotting were performed as indicated. Actin served as loading control. (n=3). (C,D and L) * indicates non-specific band.

### AMPK-dependent phosphorylation of CRY1 promotes nigericin-induced ubiquitination and degradation

Previous studies have shown that phosphorylation of CRY1 by AMPK at Ser71 facilitates its degradation via FBXL3 (17). To determine whether AMPK activation contributes to nigericin-induced CRY destabilization, we first analyzed AMPK phosphorylation in primary human macrophages. As expected, metformin (MET), used as a positive control, robustly activated AMPK, and nigericin triggered a similar phosphorylation signal (Figure 5H).

Having established that nigericin engages AMPK signaling, we next compared the kinetics of AMPK activation and CRY loss in hMDMs during nigericin treatment. The decrease in CRY abundance coincided with increased AMPK phosphorylation, supporting a link between AMPK activation and CRY destabilization in this context (Figure 5I). To directly test whether AMPK catalytic activity impacts CRY1 degradation, we used HEK293T cells expressing either wild-type AMPK (AMPK WT) or a kinase-dead mutant (AMPK KD). Following nigericin treatment, CRY1 degradation was reduced in cells expressing AMPK KD compared with AMPK WT. As phosphorylation of acetyl-CoA carboxylase 1 (ACC1), confirmed that AMPK was effectively activated in AMPK WT-expressing cells, whereas the kinase-dead mutant was unable to phosphorylate this downstream target (Figure 5J). These results indicate that AMPK enzymatic activity is important for efficient CRY1 degradation in response to nigericin. We then asked whether phosphorylation of CRY1 at the AMPK-sensitive Ser71 site is required for its ubiquitination in this setting. Using a CRY1 mutant that cannot be phosphorylated by AMPK (S71A), we found that nigericin-induced polyubiquitin chain formation on CRY1 was abolished (Figure 5K). Together with previous work identifying Ser71 as a key AMPK target site on CRY1 (35), these data support a model in which AMPK-dependent phosphorylation at Ser71 enables FBXL3-mediated CRY1 ubiquitination and degradation following nigericin stimulation.

We next asked whether AMPK-dependent phosphorylation at S71 also impacts CRY1 association with NLRP3. Under basal conditions, WT CRY1 and the S71A mutant associated similarly with NLRP3. However, after nigericin stimulation, the S71A mutant maintained stronger binding to NLRP3 (Figure 5L), consistent with impaired turnover when phosphorylation-dependent degradation is blocked. Together, these data link AMPK-dependent phosphorylation of CRY1 to FBXL3-mediated CRY ubiquitination and destabilization, and suggest that this pathway contributes to the nigericin-induced reduction of CRY-NLRP3 association.

### Domain and structural analyses define an interface between NLRP3 NACHT-LRR and the CRY PHR domain

To characterize the molecular basis of CRY-NLRP3 complex formation, we first mapped the domains responsible for CRY binding. Co-immunoprecipitation assays with truncated NLRP3 constructs in HEK293T cells indicated that the PYD domain was dispensable, whereas both the NACHT and LRR domains were capable of CRY2 association (Figure 6A, S7A). Reciprocal analysis of truncated CRY proteins showed that the photolyase homology region (PHR) was necessary for NLRP3 binding, while the C-terminal destabilization domain was not (Figure 6B, S7B), identifying the PHR domain as a conserved interface for CRYs to interact with NLRP3. Comparable mapping experiments performed with CRY1 yielded consistent results (Figure S7C). To visualize the potential structure of the human CRY2-NLRP3 complex, we used AlphaFold to model both proteins. The resulting models were analyzed with UCSF ChimeraX, and the model with the highest confidence score (model_0) was selected for further analysis (Figure 6C; Figure S7D). Inter-chain confidence was evaluated using Predicted Aligned Error (PAE) scores. The average PAE between chains A and B was approximately 30 Å, indicating flexibility in the relative orientation of the two proteins. Although overall confidence in the relative positioning of the chains was modest, the predicted structures closely matched previously published models of human NLRP3 associated with CRID3/MCC950 (PDB 7PZC; RMSD = 1.88 Å over 592 of 1010 Cα pairs; 59% sequence coverage) and mouse CRY2 associated with FBXL3 and SKP1 (PDB 4I6J; RMSD = 0.55 Å over 489/503 Cα pairs; 97% coverage) (40) (Figure S7D). We therefore used these published structures to align with the AlphaFold models and obtain higher-quality CRY2-NLRP3 and CRY1-NLPR3 complexes (Figure 6C-lower panel; Figure S7E). Consistent with this, the CRY1-NLRP3 model aligned to NLRP3 from 7PZC with an RMSD of 2.16 Å over 796 of 1010 pruned Cα pairs (79% structural coverage) (Figure S7E). PAE analysis confirmed high intra-chain confidence (minimum PAE = 0.76 Å) but low inter-chain confidence (minimum inter-chain PAE = 20 Å), consistent with flexibility while still allowing identification of amino acids likely involved at the interface. Analysis of inter-chain contacts (allowed overlap −0.1 Å; center-center distance < 2 Å) identified 36 van der Waals interactions for CRY2-NLRP3 (60 for NLRP3-CRY1), involving residues primarily localized within the NACHT and LRR domains of NLRP3 and the PHR domain of CRY2 and CRY1 (Table 1 and Table 2), supporting a structural interface in which NLRP3 engages CRYs through its NACHT-LRR region

**Figure 6.**
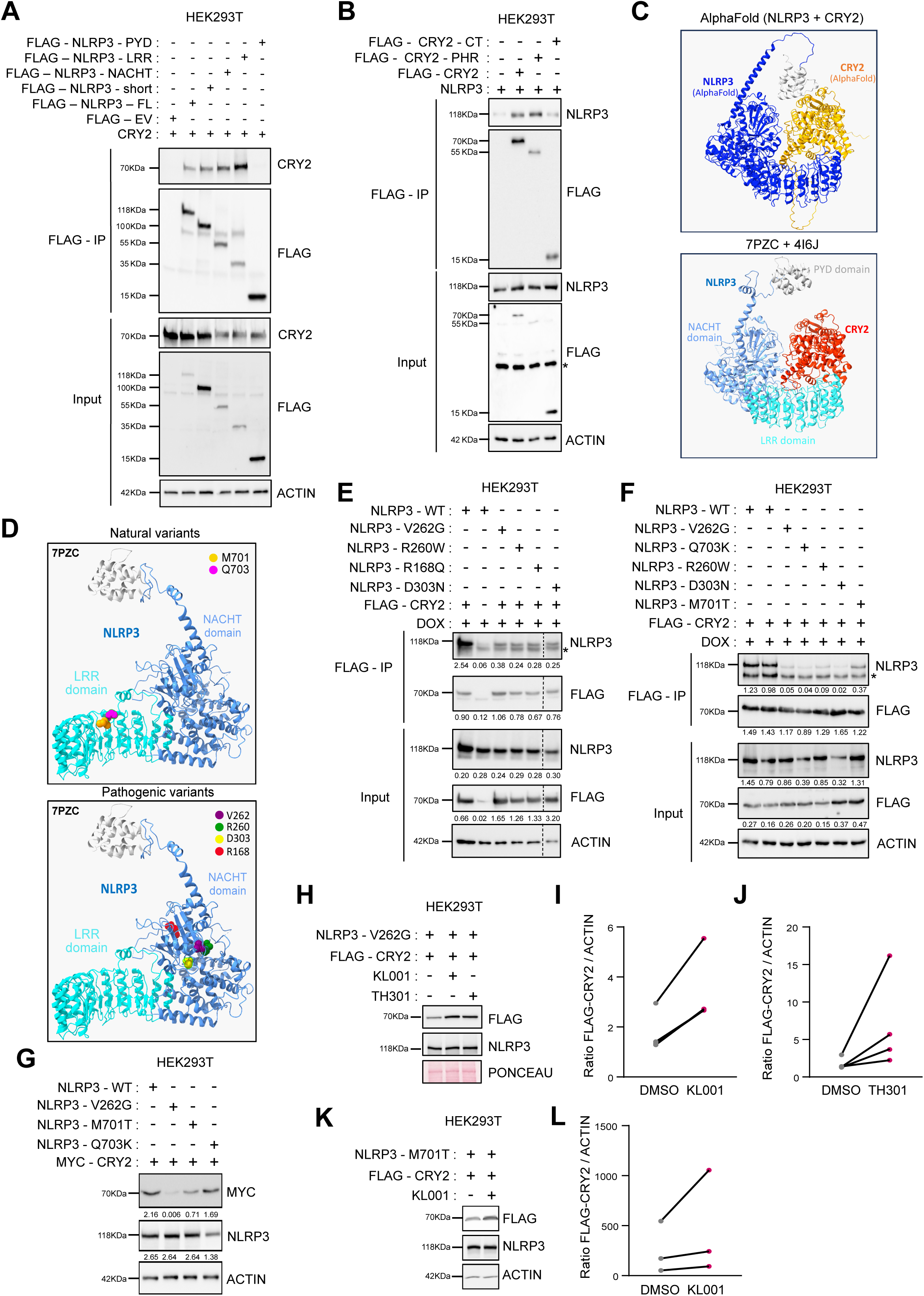
Domain and structural analyses define a CRY-NLRP3 interaction interface and identify CAPS-associated NLRP3 variants that weaken CRY binding and modulate CRY2 stability. (A) HEK293T cells were transfected with FLAG-NLRP3 constructs and CRY2 for 48 h before anti-FLAG immunoprecipitation and immunoblotting (n=3). (B) HEK293T cells were transfected with NLRP3, FLAG-CRY2-FL, or CRY2 truncations for 48 h before anti-FLAG immunoprecipitation and immunoblotting (n=3). (C) Top panel: AlphaFold-predicted model of the human CRY2-NLRP3 complex. Bottom panel: alignment of mouse CRY2 (PDB 4I6J) in red with human NLRP3 (PDB 7PZC) in blue based on the AlphaFold prediction. NLRP3 domains are color-coded as follows: PYD (gray), NACHT (dark blue), and LRR (light blue); CRY2 structure is shown in red. (D) ChimeraX structural representation of human NLRP3 showing the spatial localization of amino acid residues R168, R260, V262, D303, M701, and Q703. The PYD domain is shown in grey, NACHT and LRR domains in shades of blue. (E - F) HEK293T cells were transfected with FLAG-CRY2 and WT NLRP3 or CAPS/AID mutants (V262G, R260W, D303N, M701T, Q703K) for 48 h, followed by anti-FLAG immunoprecipitation and immunoblotting (n=4). (G) Immunoblot analysis of whole-cell lysates from HEK293T cells expressing MYC-CRY2 with WT NLRP3 or mutants V262G, M701T or Q703K. Actin served as a loading control (n=2). (H-K) HEK293T cells expressing FLAG-CRY2 and doxycycline-inducible NLRP3 V262G (H) or M701T (K) were treated with CRY stabilizers (KL001, 5 µM; TH301, 2 µM) for 24 h following doxycycline induction (1 µg/mL, 24 h) (n=3). (I, J and L) Quantification of FLAG-CRY2 protein levels from panels (H and K), normalized to actin (Image Lab). (B, E and F) * indicates non-specific band.

**Table 1:** Structural mapping of the NLRP3–CRY1 interaction interface. Residue pairs predicted to mediate the NLRP3–CRY1 interaction were identified using the NLRP3 cryo-EM structure (PDB ID: 7PZC) and the AlphaFold model of CRY1 (see Figure 6C). Amino acid positions and identities for both proteins are listed for residues located in close spatial proximity. Distances (Å) indicate the shortest atomic distances between residues. Recurrent entries correspond to multiple atomic contacts involving the same residue pair.

| PDB | aa position | aa | PDB | aa position | aa | Distance (Å) |
| --- | --- | --- | --- | --- | --- | --- |
| 7pzc | 180 | GLN | AlphaFold CRY1 model | GLU | 226 | 1.901 |
| 7pzc | 180 | GLN | AlphaFold CRY1 model | GLU | 226 | 1.645 |
| 7pzc | 180 | GLN | AlphaFold CRY1 model | GLU | 226 | 1.111 |
| 7pzc | 180 | GLN | AlphaFold CRY1 model | GLU | 226 | 1.932 |
| 7pzc | 218 | PRO | AlphaFold CRY1 model | LYS | 68 | 1.293 |
| 7pzc | 218 | PRO | AlphaFold CRY1 model | LYS | 68 | 1.049 |
| 7pzc | 218 | PRO | AlphaFold CRY1 model | LYS | 68 | 1.817 |
| 7pzc | 218 | PRO | AlphaFold CRY1 model | LYS | 68 | 1.923 |
| 7pzc | 218 | PRO | AlphaFold CRY1 model | LYS | 68 | 1.716 |
| 7pzc | 342 | PRO | AlphaFold CRY1 model | ARG | 67 | 1.791 |
| 7pzc | 360 | HIS | AlphaFold CRY1 model | SER | 207 | 1.522 |
| 7pzc | 619 | LYS | AlphaFold CRY1 model | ARG | 437 | 1.866 |
| 7pzc | 688 | PRO | AlphaFold CRY1 model | ALA | 229 | 1.590 |
| 7pzc | 688 | PRO | AlphaFold CRY1 model | ALA | 229 | 0.732 |
| 7pzc | 699 | ARG | AlphaFold CRY1 model | ARG | 238 | 1.919 |
| 7pzc | 702 | ASP | AlphaFold CRY1 model | GLU | 235 | 1.902 |
| 7pzc | 703 | MET | AlphaFold CRY1 model | GLU | 235 | 1.792 |
| 7pzc | 707 | VAL | AlphaFold CRY1 model | LEU | 225 | 1.378 |
| 7pzc | 707 | VAL | AlphaFold CRY1 model | TRP | 230 | 1.906 |
| 7pzc | 920 | ARG | AlphaFold CRY1 model | GLU | 235 | 1.854 |
| 7pzc | 1008 | MET | AlphaFold CRY1 model | LYS | 151 | 1.996 |
| 7pzc | 1008 | MET | AlphaFold CRY1 model | LYS | 151 | 1.401 |
| 7pzc | 1008 | MET | AlphaFold CRY1 model | LYS | 151 | 1.380 |
| 7pzc | 1008 | MET | AlphaFold CRY1 model | LYS | 151 | 1.263 |
| 7pzc | 1008 | MET | AlphaFold CRY1 model | LYS | 151 | 1.704 |
| 7pzc | 1008 | MET | AlphaFold CRY1 model | LYS | 151 | 1.633 |
| 7pzc | 1008 | MET | AlphaFold CRY1 model | LYS | 151 | 1.776 |
| 7pzc | 1009 | TYR | AlphaFold CRY1 model | PHE | 295 | 1.928 |
| 7pzc | 1009 | TYR | AlphaFold CRY1 model | PHE | 295 | 1.922 |
| 7pzc | 1010 | PHE | AlphaFold CRY1 model | THR | 149 | 1.831 |
| 7pzc | 1011 | ASN | AlphaFold CRY1 model | LEU | 148 | 1.950 |
| 7pzc | 1011 | ASN | AlphaFold CRY1 model | PHE | 406 | 1.870 |
| 7pzc | 1012 | TYR | AlphaFold CRY1 model | PHE | 406 | 1.918 |
| 7pzc | 1012 | TYR | AlphaFold CRY1 model | PHE | 406 | 1.321 |
| 7pzc | 1012 | TYR | AlphaFold CRY1 model | TYR | 488 | 1.958 |
| 7pzc | 1012 | TYR | AlphaFold CRY1 model | TYR | 488 | 1.695 |
| 7pzc | 1018 | LEU | AlphaFold CRY1 model | GLN | 408 | 1.621 |
| 7pzc | 1018 | LEU | AlphaFold CRY1 model | GLN | 408 | 1.835 |
| 7pzc | 1018 | LEU | AlphaFold CRY1 model | GLN | 408 | 1.147 |
| 7pzc | 1018 | LEU | AlphaFold CRY1 model | GLN | 408 | 1.982 |
| 7pzc | 1019 | GLU | AlphaFold CRY1 model | PHE | 410 | 1.993 |
| 7pzc | 1023 | GLU | AlphaFold CRY1 model | PHE | 410 | 1.543 |
| 7pzc | 1023 | GLU | AlphaFold CRY1 model | PHE | 410 | 1.635 |
| 7pzc | 1036 | TRP | AlphaFold CRY1 model | TRP | 399 | 1.716 |
| 7pzc | 1036 | TRP | AlphaFold CRY1 model | TRP | 399 | 1.437 |
| 7pzc | 1036 | TRP | AlphaFold CRY1 model | TRP | 399 | 1.460 |
| 7pzc | 1036 | TRP | AlphaFold CRY1 model | TRP | 399 | 1.736 |
| 7pzc | 1036 | TRP | AlphaFold CRY1 model | GLN | 407 | 1.873 |
| 7pzc | 1036 | TRP | AlphaFold CRY1 model | TRP | 399 | 1.550 |
| 7pzc | 1036 | TRP | AlphaFold CRY1 model | TRP | 399 | 1.422 |
| 7pzc | 1036 | TRP | AlphaFold CRY1 model | TRP | 399 | 1.773 |
| 7pzc | 1036 | TRP | AlphaFold CRY1 model | TRP | 399 | 1.952 |
| 7pzc | 1036 | TRP | AlphaFold CRY1 model | TRP | 399 | 1.415 |
| 7pzc | 1036 | TRP | AlphaFold CRY1 model | TRP | 399 | 1.622 |
| 7pzc | 1036 | TRP | AlphaFold CRY1 model | TRP | 399 | 1.783 |
| 7pzc | 1036 | TRP | AlphaFold CRY1 model | TRP | 399 | 1.483 |
| 7pzc | 1036 | TRP | AlphaFold CRY1 model | TRP | 399 | 1.804 |
| 7pzc | 1036 | TRP | AlphaFold CRY1 model | TRP | 399 | 1.206 |
| 7pzc | 1036 | TRP | AlphaFold CRY1 model | TRP | 399 | 1.225 |
| 7pzc | 1036 | TRP | AlphaFold CRY1 model | TRP | 399 | 1.521 |

**Table 2:** Predicted NLRP3–CRY2 contact residues identified from structural docking. Residue pairs predicted to contribute to the NLRP3–CRY2 interaction were identified using the NLRP3 cryo-EM structure (PDB ID: 7PZC) and the CRY2 crystal structure (PDB ID: 4I6J) (see Figure 6C). Amino acid positions and identities are listed for residues positioned in close spatial proximity. Distances (Å) indicate the shortest atomic distances calculated between residues. Recurrent entries reflect multiple atomic contacts involving the same residue pair.

| PDB | aa position | aa | PDB | aa position | aa | Distance (Å) |
| --- | --- | --- | --- | --- | --- | --- |
| 7pzc | 100 | GLU | 4i6j | 428 | PHE | 1.960 |
| 7pzc | 180 | GLN | 4i6j | 508 | GLN | 1.590 |
| 7pzc | 180 | GLN | 4i6j | 508 | GLN | 1.818 |
| 7pzc | 180 | GLN | 4i6j | 508 | GLN | 1.953 |
| 7pzc | 180 | GLN | 4i6j | 508 | GLN | 1.446 |
| 7pzc | 180 | GLN | 4i6j | 508 | GLN | 1.213 |
| 7pzc | 363 | ASP | 4i6j | 328 | GLU | 1.789 |
| 7pzc | 363 | ASP | 4i6j | 328 | GLU | 0.972 |
| 7pzc | 363 | ASP | 4i6j | 328 | GLU | 1.941 |
| 7pzc | 363 | ASP | 4i6j | 328 | GLU | 1.974 |
| 7pzc | 685 | HIS | 4i6j | 511 | ARG | 1.039 |
| 7pzc | 688 | PRO | 4i6j | 512 | TYR | 1.705 |
| 7pzc | 688 | PRO | 4i6j | 512 | TYR | 1.006 |
| 7pzc | 689 | LYS | 4i6j | 512 | TYR | 1.624 |
| 7pzc | 689 | LYS | 4i6j | 512 | TYR | 1.891 |
| 7pzc | 703 | MET | 4i6j | 517 | LEU | 1.139 |
| 7pzc | 703 | MET | 4i6j | 517 | LEU | 1.811 |
| 7pzc | 703 | MET | 4i6j | 517 | LEU | 1.348 |
| 7pzc | 703 | MET | 4i6j | 517 | LEU | 1.704 |
| 7pzc | 704 | VAL | 4i6j | 520 | SER | 1.491 |
| 7pzc | 705 | GLN | 4i6j | 503 | LYS | 1.509 |
| 7pzc | 705 | GLN | 4i6j | 503 | LYS | 1.477 |
| 7pzc | 705 | GLN | 4i6j | 503 | LYS | 1.487 |
| 7pzc | 706 | CYS | 4i6j | 507 | GLN | 1.466 |
| 7pzc | 706 | CYS | 4i6j | 507 | GLN | 1.367 |
| 7pzc | 707 | VAL | 4i6j | 505 | ILE | 1.389 |
| 7pzc | 779 | ARG | 4i6j | 513 | ARG | 1.672 |
| 7pzc | 779 | ARG | 4i6j | 513 | ARG | 1.802 |
| 7pzc | 779 | ARG | 4i6j | 513 | ARG | 1.787 |
| 7pzc | 779 | ARG | 4i6j | 513 | ARG | 1.897 |
| 7pzc | 835 | VAL | 4i6j | 515 | LEU | 1.384 |
| 7pzc | 835 | VAL | 4i6j | 515 | LEU | 0.985 |
| 7pzc | 835 | VAL | 4i6j | 515 | LEU | 1.404 |
| 7pzc | 1007 | GLU | 4i6j | 428 | PHE | 1.949 |
| 7pzc | 1007 | GLU | 4i6j | 429 | HIS | 1.856 |
| 7pzc | 1012 | TYR | 4i6j | 256 | ARG | 1.648 |

### NLRP3 variants at the predicted CRY interface and pathogenic CAPS mutations weaken CRY binding and differentially affect CRY2 stability

Analysis of the predicted CRY-NLRP3 interface revealed a set of NLRP3 residues predicted to contribute to CRY binding (Table 1 and Table 2). Notably, the shortlist highlighted LRR-domain positions corresponding to variants reported in patients diagnosed with Cryopyrin-Associated Periodic Syndromes (CAPS), a group of rare monogenic autoinflammatory diseases caused by gain-of-function mutations in NLRP3 that drive excessive IL-1β production and systemic inflammation (41), including M701T and Q703K (Figure 6D, upper). Because the pathogenicity of M701T and Q703K remains debated or not fully established, we refer to these as NLRP3 interface variants. Clinically, CAPS encompass a spectrum of disorders historically classified according to severity as familial cold autoinflammatory syndrome (FCAS), Muckle-Wells syndrome (MWS), and neonatal-onset multisystem inflammatory disease (NOMID) (42,43). Many established pathogenic CAPS gain-of-function mutations cluster within the NACHT domain and destabilize the inactive conformation of NLRP3. Because our domain mapping and structural models indicate that CRY engages NLRP3 through its NACHT-LRR region, we reasoned that NACHT mutations that favor an “open/active” state could weaken CRY binding by altering the CRY-interaction surface. We therefore selected representative pathogenic NACHT mutations (R168Q, R260W, V262G, and D303N), which map to the NBD/HD1 subdomains near the nucleotide-binding pocket and have been implicated in maintaining the inactive state (Figure 6D, lower). Because inflammasome activation is accompanied by loss of CRY-NLRP3 complexes, we further hypothesized that constitutively activating NACHT-domain CAPS mutations, as well as LRR NLRP3 interface variants, might weaken CRY binding and thereby impact CRY stability. We therefore compared in parallel (i) LRR NLRP3 interface variants highlighted by the structural predictions and (ii) established pathogenic NACHT-domain CAPS gain-of-function mutations.

To test whether NLRP3 variants alter CRY association, we co-expressed FLAG-CRY2 with WT NLRP3 or the indicated NLRP3 variants in HEK293T cells and assessed binding by co-immunoprecipitation (Figure 6E,F). FLAG-CRY2 inputs were normalized by adjusting protein amounts, ensuring that differences in co-precipitation reflected NLRP3 variation rather than CRY expression levels. Across the panel, NLRP3 variants reduced CRY2 binding compared with WT NLRP3 (Figure 6E,F). Similar results were obtained with CRY1 (Figure S7F, G).

We next asked whether the reduced CRY-NLRP3 association observed for these variants was accompanied by changes in CRY2 abundance. Consistent with the decrease in CRY abundance observed upon inflammasome activation, several NLRP3 variants were associated with reduced CRY2 levels. Immunoblot analysis of CRY2 expression showed mutation-dependent effects, with the most pronounced decrease observed for the pathogenic CAPS mutant V262G, a similarly strong reduction for M701T, and a more modest effect for Q703K (Figure 6G). Given the strong impact of V262G on CRY2 abundance, we focused on this variant to test whether pharmacological stabilization could restore CRY2 levels. Treatment with the CRY stabilizers KL001 or TH301 increased CRY2 abundance in the presence of V262G (Figure 6H-J). KL001 modestly and similarly increased CRY2 abundance in the context of M701T mutant (Figure 6K-L). Together, these experiments show that both interface-positioned variants and pathogenic NACHT mutations reduce CRY-NLRP3 interaction, while effects on CRY2 abundance are variant-dependent, consistent with potentially distinct underlying molecular mechanisms.

### Time after synchronization modulates MCC950 efficacy, and CAPS-associated NLRP3 variants show distinct timing of maximal inhibition

Pharmacological inhibition of NLRP3 has strong therapeutic potential, given that aberrant NLRP3 signaling contributes to CAPS and to a broader spectrum of common inflammatory and degenerative pathologies. MCC950 is the best characterized NLRP3 inhibitor currently available and blocks NLRP3 activation by binding the NACHT region and stabilizing an inactive state (39,44). Because we show that CRY-NLRP3 binding oscillates over time after synchronization and inversely correlates with inflammasome outputs, we asked whether MCC950 efficacy is similarly modulated by time after synchronization, and whether such modulation is altered by NLRP3 variants tested in CAPS-associated contexts.

In synchronized hMDMs, cells were treated with nigericin in the presence or absence of MCC950 pretreatment (Figure 7A; Figure S8A). MCC950 significantly reduced cell death as measured by DRAQ7 incorporation (Figure 7A). Quantification of DRAQ7 area under the curve (AUC) showed that cell death varied with time after synchronization (Figure 7B). Both time after synchronization and MCC950 treatment significantly affected this readout, with a trend toward stronger protection at 30 h than at 18 h after synchronization. However, we did not detect a significant interaction between time and MCC950 in this dataset, limiting our ability to draw firm conclusion about time-dependent differences in MCC950 efficacy based on cell death alone (Figure 7A,B).

**Figure 7.**
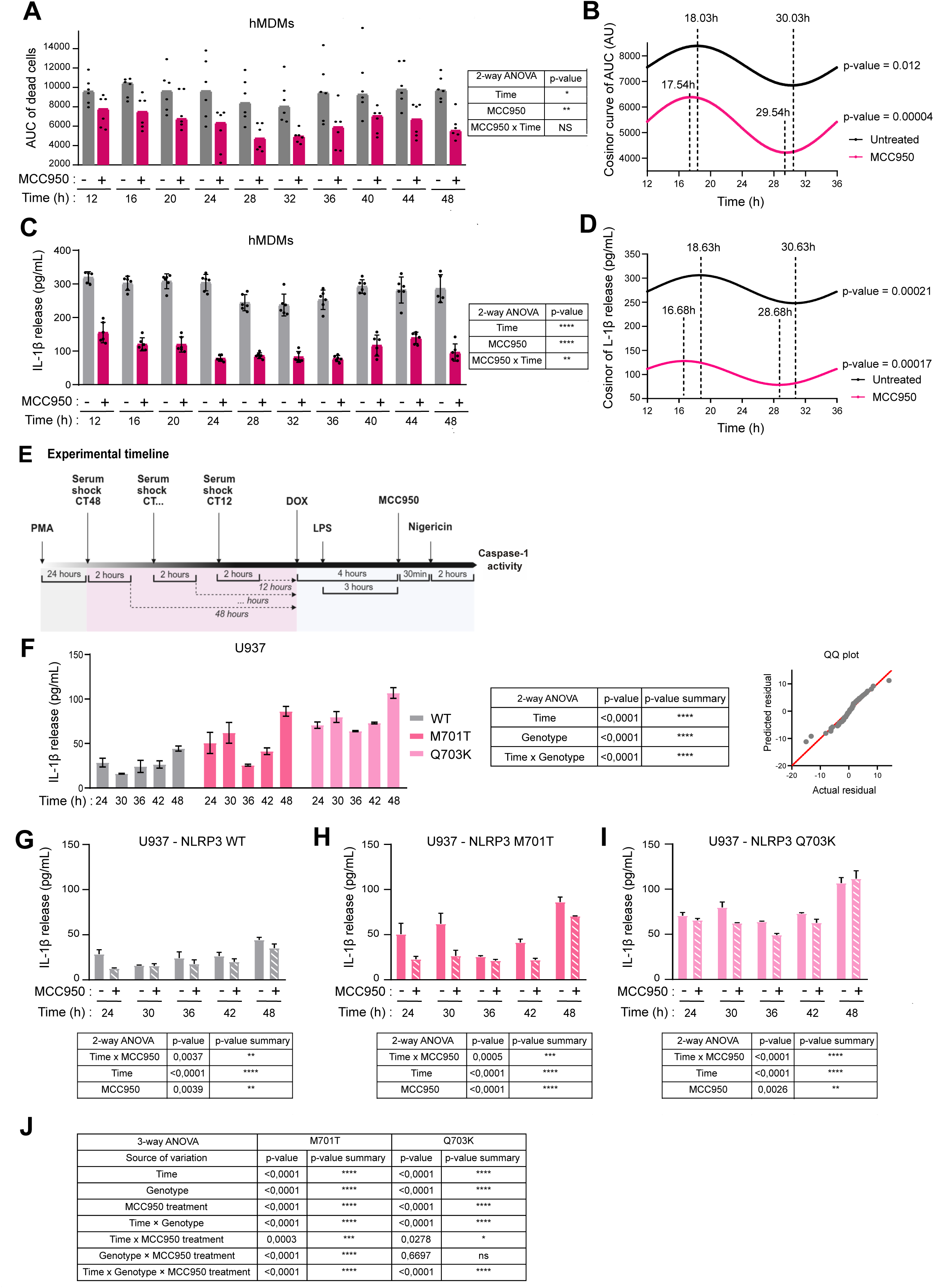
Time after synchronization modulates MCC950 efficacy in primary human macrophages and U937 cells expressing NLRP3 variants. (A) hMDMs were synchronized as in Figure S5, primed with LPS (0.5 µg/mL, 4 h), and treated or not with MCC950 (150 nM, 30 min pretreatment) before stimulation with nigericin (20 µM). Cell death was monitored over time by DRAQ7 incorporation using an Opera Phenix HCS microscope. Area under the curve (AUC) quantification of cell-death time courses from (Figure S8A). Data represent n = 4 independent experiments with 6 replicates each; two-way repeated-measures ANOVA with time and treatment as factors and individual donors, ns = not significant, *p < 0.03, ***p = 0.0002. (B) Cosinor analysis of AUC data from (A), showing acrophase, bathyphase, and p-values. (C) IL-1β secretion measured by ELISA in supernatants from hMDMs at each time point after synchronization. The untreated condition corresponds to the same samples shown in Figure 5L and is displayed here for direct comparison with MCC950-treated cells. Data are from three independent experiments, each performed with two experimental replicates (6 wells per condition), shown as mean ± SD. (D) Cosinor analysis of IL-1β secretion data from (C), showing acrophase, bathyphase, and p-values. (E) Experimental timeline for U937 cells expressing doxycycline-inducible NLRP3 WT or CAPS variants used in panels (F-J). (F-I) IL-1β secretion measured by ELISA in culture supernatants from synchronized U937 cells collected at 24 h, 30 h, 36 h, 42 h, and 48 h post-synchronization. Cells expressing WT NLRP3 (F, G), M701T (F, H), or Q703K (F, I) were treated or not with MCC950. (F) Left panel: IL-1β quantification across time. Middle panel: summary table reporting two-way ANOVA results. Right panel: Q-Q plot assessing normality of the IL-1β data. (G-I) IL-1β secretion profiles for each NLRP3 genotype treated or left untreated with MCC950, along with the corresponding two-way ANOVA tables. (J) Summary tables of three-way ANOVA results for panels (G-I).

We therefore examined inflammasome output using IL-1β secretion. In synchronized hMDMs, nigericin-induced IL-1β release varied over time after synchronization and was strongly reduced by MCC950 (Figure 7C-D). Notably, the magnitude of inhibition differed across timepoints, with stronger suppression at 24 h after synchronization and reduced efficacy at 12 h and 44 h after synchronization. Cosinor fitting supported significant rhythmicity of IL-1β secretion in both conditions, with overall lower IL-1β levels and a shift in the timing of peak secretion upon MCC950 treatment (Figure 7C). Together, these data indicate that inflammasome output and its pharmacological inhibition by MCC950 are modulated by time after synchronization in primary human macrophages.

We next asked whether time-dependent regulation of inflammasome activity and MCC950 efficacy extends to NLRP3 genetic variation relevant to CAPS cohorts. To address this, we used U937 cells carrying doxycycline-inducible WT NLRP3 or NLRP3 variants, which were differentiated and synchronized using a protocol comparable to that used for hMDMs. Cells were then treated with doxycycline for 4 h to induce NLRP3 expression, primed with LPS for 3 h, and treated or not with MCC950 for 30 min before nigericin stimulation, following a defined timeline (Figure 7E). Before testing MCC950, we assessed time-dependent inflammasome outputs across genotypes. In WT U937 cells, both IL-1β secretion (Figure 7F) and caspase-1 activity (Figure S8B-E; G-I; L,N) varied reproducibly over time after synchronization. We then compared (i) the established pathogenic CAPS gain-of-function mutant V262G and (ii) variants reported in CAPS cohorts with unestablished pathogenicity, including M701T and Q703K. In this experimental system, M701T and Q703K were associated with elevated IL-1β secretion, with variant- and time-dependent differences in amplitude and timing compared to WT cells (Figure 7F; S8D).

We then assessed how NLRP3 variants influences the timing of MCC950-mediated inhibition. In WT U937 cells, MCC950 robustly suppressed IL-1β secretion and caspase-1 activity, with the degree of inhibition varying over time after synchronization, mirroring the time-dependent efficacy observed in hMDMs (Figure 7G, S8E; I; L;N). In cells expressing M701T or Q703K, MCC950 also significantly reduced IL-1β secretion; however, the timepoint at which inhibition was maximal differed between variants (Figure 7H-J). At the level of caspase-1 activity, MCC950 reduced inflammasome activation across all genotypes (Figure S8E-L). However, the timing and magnitude of this inhibition varied between NLRP3 variants: in some mutants, caspase-1 activity was most strongly suppressed at earlier time points after synchronization, whereas in others maximal inhibition occurred later. Together, these results demonstrate that time after synchronization modulates both NLRP3 inflammasome activity and MCC950 efficacy, with different variants altering the timing of maximal inhibition while preserving overall sensitivity to the drug.

## Discussion

This study identifies CRY-NLRP3 complexes as a protein-level mechanism integrating circadian timing with inflammasome activation. While circadian influences on innate immunity have largely been attributed to transcriptional and metabolic programs (33–35), our findings reveal that clock components modulate inflammasome assembly through dynamic protein-protein interactions. CRY1 and CRY2 associate with NLRP3 and restrain its activation in human macrophages, establishing a temporally regulated checkpoint operating at the level of the inflammasome complex itself. Although our data do not establish direct binding in vitro, the consistent association observed by co-immunoprecipitation and proximity ligation assays supports a functional interaction that shapes inflammasome responsiveness

Our data indicate that the temporal dynamics of CRY-NLRP3 association cannot be explained solely by oscillations in CRY abundance. Although total CRY levels inversely correlate with inflammasome activity, peak CRY-NLRP3 association does not strictly coincide with maximal CRY expression, and CRY stabilization dampens inflammasome outputs without altering NLRP3 levels. These observations suggest that regulated complex formation and dissociation, rather than expression alone, contribute to temporal modulation of inflammasome activity.

Upon inflammasome stimulation, CRY-NLRP3 complexes dissociate and CRY abundance declines. However, the causal relationship between these events remains unresolved. It is unclear whether CRY degradation-potentially mediated by FBXL3-is required to release NLRP3 from inhibitory binding, or whether complex dissociation precedes and allows CRY destabilization. The observation that certain NLRP3 variants lose CRY binding without uniformly triggering CRY degradation argues against CRY destabilization being strictly required for inflammasome activation. Whether ionic imbalance directly influences CRY stability, FBXL3 engagement, or higher-order inflammasome assembly will require further investigation.

Our findings implicate AMPK as an upstream regulator of CRY1 turnover in this context. Nigericin perturbs ion homeostasis and mitochondrial function, conditions consistent with AMPK activation. Although AMPK is frequently described as a negative regulator of inflammasome signaling, particularly under metabolic stress where it limits mitochondrial ROS production, promotes mitophagy, and dampens NF-κB-dependent priming (45,46), its effects are highly context dependent. In human macrophages, nigericin rapidly induces AMPK phosphorylation and is accompanied by a reduction in CRY protein abundance. Consistent with this model, kinase-dead AMPK and CRY1 S71A mutants attenuate nigericin-induced CRY1 degradation, supporting a role for AMPK activity in FBXL3-mediated CRY1 destabilization. However, no direct AMPK phosphorylation site has yet been identified on CRY2, and further investigation will be required.

We next examined how NLRP3 variants reported in CAPS-associated contexts intersect with the CRY-NLRP3 regulatory interface. Mapping these mutations onto our structural model revealed that both established gain-of-function NACHT mutations and LRR variants localize near the predicted CRY-contact surface. Consistent with this positioning, all mutants tested displayed reduced CRY-NLRP3 association relative to wild-type NLRP3. However, no clear relationship emerged between the extent of CRY-NLRP3 interaction loss and reported disease severity. Nevertheless, our data further suggest that CRY stability is influenced, but not uniformly dictated, by this interaction. In our working model, NLRP3 binding shields CRY from ubiquitin-dependent degradation, such that weakened association would increase CRY susceptibility to turnover. Several NACHT mutations and related variants fit this model, showing both reduced CRY binding and enhanced CRY destabilization. Yet other mutants with comparable reductions in binding had only modest effects on CRY abundance. Thus, while impaired CRY association appears to be a shared feature of these variants, its impact on CRY stability is mutation-specific. These findings imply that additional factors-such as differential engagement of ubiquitin ligases or other CRY-associated regulators shape CRY turnover in ways that remain to be clarified.

Pharmacological inhibition of NLRP3 is being actively pursued for the treatment of inflammatory and autoinflammatory diseases. MCC950, the best-characterized inhibitor in this class, stabilizes an inactive conformation of the NACHT domain (39,44). Our data indicate that MCC950 efficacy is not temporally uniform but varies across the circadian cycle in primary human macrophages. These findings identify circadian timing as a determinant of pharmacological responsiveness and suggest that drug efficacy may depend, at least in part, on the temporal state of the inflammasome machinery.

In CAPS-associated contexts, this temporal dimension becomes mutation-specific. Although all NLRP3 variants examined remained responsive to MCC950, the phase of maximal inhibition differed between genotypes. Thus, the efficacy of NLRP3 inhibition depends on both circadian timing and mutation class. While in vivo chronotherapy studies will be required to evaluate this concept clinically, our findings provide a mechanistic rationale for integrating temporal context and genetic background when optimizing NLRP3-targeted therapies.

Recent work has demonstrated that CRY stability can be pharmacologically targeted in human disease models. In cancer settings, CRY-stabilizing compounds reduce proliferation and alter survival pathways, and isoform-selective CRY activators show translational promise (47,48). Stabilization of CRY proteins produces measurable therapeutic effects in these systems, and our findings extend this concept to innate immunity by showing that CRY stabilization dampens NLRP3 inflammasome activation in primary human macrophages. Together, these observations identify CRY proteins as post-translational regulators of NLRP3 inflammasome activity and support a model in which CRY stability influences inflammatory output beyond previously described transcriptional and metabolic circadian pathways.

## Materiel and Methods

### Cell Culture

U2OS, HEK293T, and THP-1 cells were obtained from ATCC, and THP-1 KO cells were from Invivogen. U2OS and HEK293T cells were cultured in DMEM (Gibco, 41965-039) supplemented with 10% fetal bovine serum (FBS; Gibco, 10270-106), 1% penicillin-streptomycin (Invitrogen, 15140122), 1% GlutaMAX (Invitrogen, 35050061), and 1% sodium pyruvate (Invitrogen, 11360070). THP-1 and THP-1 KO cells were maintained in RPMI 1640 (Invitrogen, 21875034) with 10% FBS, 1% penicillin-streptomycin, 1% GlutaMAX, 1% sodium pyruvate, and 50 μM β-mercaptoethanol (Sigma-Aldrich, M3148). U937 cells were cultured in RPMI 1640 with 10% FBS, 1% penicillin-streptomycin, and 1% GlutaMAX. THP-1 monocytes were differentiated into macrophages using 0.5 μL/mL phorbol 12-myristate 13-acetate (PMA; Sigma-Aldrich, P-1585) for 3 h at 37°C, followed by washing and resuspension in fresh medium. U937 monocytes were differentiated with 40 ng/mL PMA for 20 h at 37°C before medium replacement. All cell lines were maintained at 37°C in 5% CO₂ and tested negative for mycoplasma contamination (Lonza, LT07-318).

### Generation of Human Monocyte-Derived Macrophages (hMDMs)

Human peripheral blood mononuclear cells were isolated from healthy donor blood (EFS, Lyon, France) by density centrifugation using Ficoll (Cytiva, 17144003). Monocytes were purified using Percoll (EuroBio, CMSMSL01-01) and differentiated into macrophages by culture for 7 days in RPMI 1640 supplemented with 10% FBS, 1% penicillin-streptomycin, 1% GlutaMAX, and 50 ng/mL M-CSF (Proteintech, HZ-1192). Medium was refreshed on days 3 and 6.

Macrophage purity was assessed by flow cytometry using the following antibodies: CD14 (M5E2), CD1a (HI149), CD86 (2331, FUN-1), CD206 (15–02), CD163 (GHI/61), CD3 (UCHT1), CD15 (MC480), CD19 (HIB19), CD56 (NCAM16.2), CD64 (10.6), and HLA-DR (G46-6) (BD Biosciences/eBiosciences). Cells were stained after FcR blocking (Miltenyi Biotec) and analyzed on a NovoCyte Opteon flow cytometer (Agilent). Data were processed using FlowJo v10.10 (Tree Star Inc).

### Cell Synchronization

Cells were synchronized by incubation in medium containing 70% FBS for 2 h at 37°C, followed by replacement with normal medium to define circadian time 0 (CT0). Cells were harvested every 4 h from CT12 to CT48, or sequentially synchronized every 4 h and analyzed together at CT0.

### Transfection

HEK293T cells were plated at 2×10⁶ cells per 10-cm dish and transfected with 3 μg total plasmid DNA using polyethylenimine (PEI; Polyscience Inc., 23966-2) according to the manufacturer’s protocol. DNA and PEI were mixed in DMEM (1 mL) at a 1:20 ratio (DNA:PEI, μg:μL), incubated 20 min at room temperature, and added dropwise to cells. Medium was replaced after 24 h, and experiments were performed 36 h post-transfection.

### Plasmids

The following plasmids were used for transfection: pCR3-FLAG-NLRP3, pCR3-FLAG-empty, pCR3-FLAG-NLRP3-NACHT, pCR3-FLAG-NLRP3-LRR, pCR3-FLAG-NLRP3-PYD, pCR3-FLAG-NLRP3-ΔPYD, FLAG-ASC (gifts from the Tschopp laboratory); pcDNA3-NLRP3, pINDUCER21-NLRP3-V262G, pINDUCER21-NLRP3-R260W, pINDUCER21- NLRP3-R168Q, pINDUCER21-NLRP3-D303N (gifts from the Bénédicte Py laboratory); pcDNA3-empty (gift from Kyle Miller laboratory); pcDNA3-HA-empty, pBABE-CRY1, pBABE-CRY2, pcDNA3-HA-CRY1, pcDNA3-HA-CRY2, pcDNA3-FLAG-REV-ERBα, pcDNA3-PER2-V5, pcDNA3-CLOCK-HA, pcDNA3-BMAL1-MYC, pcDNA3-NPAS2-MYC, pcDNA3-CRY1-MYC, pcDNA3-CRY2-MYC, pCR3-FLAG-CRY2-PHR, pCR3-FLAG-CRY2-CT, pLenti-V5-FBXL3, pcDNA3-HA-CRY2-I499D, pcDNA3-HA-CRY2- F428D, pcDNA3-HA-CRY2-L517D, and HA-Ubiquitin (gifts from the Katja Lamia laboratory).

### CRY Stabilizer Treatment

Cells were pretreated for 48 h with KL001 (5 μM; TargetMol, T22891), KL044 (5 μM; TargetMol, #1801856-93-8), or TH301 (1-2 μM; TargetMol). DMSO served as vehicle control.

### Inflammasome Activation

THP-1 and hMDMs were primed with LPS (0.5 μg/mL; InvivoGen, tlrl-3pelps) for 3 h; U937 cells were primed with LPS (40 ng/mL) in RPMI 1640. Medium was then replaced with Opti-MEM (Gibco, 11058-021) containing 10 μM nigericin (InvivoGen, tlrl-nig) or vehicle for 30 min-3 h as indicated.

For inhibition experiments, cells were pretreated with 150 nM MCC950 (Adipogen, AG-CR1-3615) for 30 min before stimulation.

### Protein Extraction and Western Blotting

Cells were lysed in 2× Laemmli buffer (0.125 M Tris-HCl, pH 6.8; 2% SDS; 0.5 M DTT) and protein concentrations quantified by Bradford assay (Bio-Rad, 5000006) using a BSA standard curve (Sigma-Aldrich, A9647). Equal protein amounts were denatured in 6× Laemmli buffer at 95°C for 5 min, resolved by SDS-PAGE, and transferred to nitrocellulose membranes (Bio-Rad, 17001918). Membranes were blocked in 5% milk in T-TBS (20 mM Tris, 150 mM NaCl, 0.1% Tween-20) for 1 h, probed with primary antibodies overnight at 4°C. Primary antibodies included: anti-NLRP3/NALP3 (Cryo-2, Adipogen, AG-20B-0014-C100), anti-CRY1 (ABClonal, A18028), anti-CRY2 (Bethyl, A302-615A), anti-Actin (MP Biomedicals, 0869100-CF), anti-Rev-Erbα (Cell Signaling, 13418), anti-FLAG (Sigma-Aldrich, F7425), anti-V5-Tag (Cell Signaling, 13202), anti-α-Tubulin (Sigma-Aldrich, T5168), anti-HA (Sigma-Aldrich, H6908), anti-Myc-Tag (Cell Signaling, 2276), anti-GSDMD (Abcam, ab209845; Sigma-Aldrich, G7422), anti- IL-1β (R&D Systems, AF-401-NA; Cell Signaling, 12242), anti-ASC (Adipogen, AG-25B-0006), anti-Caspase-1 (Bally-1, Adipogen, AG-20B-0048), anti-P-AMPK (Thr172, Cell Signaling, 2535T), anti-AMPK (Cell Signaling, 2532S) and anti-P-ACC1 (Cell signaling, 3662S). The membranes were washed, and incubated with HRP-conjugated secondary antibodies (Promega, W402B, W401B) for 1 h at room temperature. Signals were developed using Clarity™ ECL (Bio-Rad, 170-5061) and imaged on a ChemiDoc Imaging System (Bio-Rad). Quantification was performed with Image Lab software (Bio-Rad).

### Co-Immunoprecipitation

HEK293T cells were lysed in IP buffer (50 mM HEPES pH 7.4, 138 mM KCl, 4 mM NaCl, 50 mM sodium pyrophosphate, 100 mM sodium fluoride, 10 mM EDTA, 0.1% Triton X-100, 1 mM EGTA, 1 mM phenylmethylsulfonyl fluoride, protease inhibitors) for 20 min at 4°C. Lysates were clarified by centrifugation (13,200 rpm, 30 min, 4°C) and normalized by Bradford assay. Immunoprecipitation was performed overnight at 4°C using anti-FLAG (Sigma-Aldrich, A2220) or anti-HA (Sigma-Aldrich, A2095) agarose beads. Beads were washed, boiled in Laemmli buffer, and analyzed by Western blotting.

### ELISA

IL-1β levels in cell culture supernatants were quantified using a sandwich ELISA based on the DuoSet Human IL-1β/IL-1F2 Development System (R&D Systems), following the manufacturer’s recommendations. Samples and standards were assayed in duplicate. Color development was quantified by measuring absorbance using a TECAN microplate reader (Phoenix Research Products)

### Proximity Ligation Assay (PLA)

hMDMs, THP-1, and U937 cells were seeded in 96-well plates (PhenoPlate-96, Revvity, 6055300) at 40,000 (hMDMs) or 60,000 (THP-1 or U937) cells per well. After circadian synchronization or NLRP3 inflammasome activation, cells were washed with cold DPBS and fixed in 4% paraformaldehyde diluted in DPBS (Electron Microscopy Sciences, 15710) for 15 min at room temperature. Cells were washed twice with DPBS and permeabilized with 100% cold methanol (VWR, 20847.307) for 5 min. PLA was performed using the Duolink In Situ kit (Sigma-Aldrich) according to the manufacturer’s instructions. Cells were blocked for 1 h in Duolink blocking solution and incubated for 1 h with the following primary antibody pairs: mouse anti-NLRP3 (Cryo-2, Adipogen, AG-20B-0014-C100) with either rabbit anti-CRY1 (ABClonal, A18028; ABClonal, A13666) or rabbit anti-CRY2 (Bethyl, A302-615A) or mouse anti-ATM ((2C1(1A1)) ab78) with rabbit anti-ATM ((Y170) ab32420). Samples were then incubated with PLA Probe Anti-Mouse MINUS (Sigma-Aldrich, DUO92004) and PLA Probe Anti-Rabbit PLUS (Sigma-Aldrich, DUO92002) for 1 h. Ligation (30 min) and amplification (100 min; Far Red reagents, DUO92013) were performed at 37°C in a humidified chamber with washes in Duolink wash buffer (DUO82049) between steps. After staining, wells were filled with 1× DPBS, sealed with parafilm, and stored at 4°C until imaging. Nuclei were counterstained with Hoechst (Invitrogen, H3570) for 30 min prior to image acquisition using a Revvity Opera Phenix Plus microscope (Revvity) equipped with a 60× water-immersion objective at the Cell Imaging Platform of the CRCL (PIC, CRCL, Lyon). The total PLA signal area per condition was quantified using Harmony software (Revvity), with 500–3,000 cells analyzed per condition.

### Cell Death Assay

hMDMs were treated with LPS (0.5 μg/mL, 4 h); U937 cells were treated with doxycycline (1 μg/mL, 3 h) then LPS (40 ng/mL, 2 h). After priming, medium was replaced with Opti-MEM containing DRAQ7 (1:1000; BioLegend, 424001) and Hoechst 33342 (1:100,000; Invitrogen, H3570) for 30 min. Cells were stimulated with Nigericin (20μM) during live imaging on the Opera Phenix microscope. Images were acquired every 6-10 min (20× air objective), and cell death quantified with Harmony software.

### ASC Speck Formation

LPS-primed cells were activated with NLRP3 agonists, fixed in 4% paraformaldehyde diluted in DPBS (20 min), quenched with 0.1 M glycine (10 min), permeabilized with 0.1% Triton X-100/5% BSA (1 h), and stained with anti-ASC (1:1000; rabbit, AG-25B-0006) overnight at 4°C, followed by Alexa Fluor 488 anti-rabbit IgG (1:200; Invitrogen, A-11034). Nuclei were counterstained with Hoechst. Images were acquired on the Opera Phenix HCS microscope (Revvity, 40× water objective) and ASC speck-positive cells quantified using Opera Phenix/Harmony software.

### Caspase-1 Activity

LPS-primed macrophages were treated with vehicle or nigericin for 1-3 h. Caspase-1 activity was measured using the Caspase-Glo® 1 Inflammasome Assay (Promega, G9951) following manufacturer’s instructions. Luminescence was recorded using a TECAN microplate reader and normalized to cell density quantified by crystal violet staining (30% acetic acid, 10% methanol; absorbance 595 nm).

### Protein Structure Prediction and Structural Alignment

Protein structures were predicted using the AlphaFold Server (AlphaFold v3.0) (49) on September 8, 2025. Sequences NP_004886.3 (NLRP3) and NP_066940.3 (CRY2) were truncated to residues 1-517. The rank_0 model was selected among five ranked predictions. pLDDT and PAE scores were used for model evaluation. Structures were visualized and aligned in UCSF ChimeraX (v1.10.1) (50–52) with MatchMaker (default settings) using PDB 4I6J, 6NPY, and 7PZC as references. RMSD values were computed and structures rendered for figure preparation. Van der Waals contacts between NLRP3 and CRY2 were identified in UCSF ChimeraX using the “Find Contacts” tool, restricting the analysis to non-bonded heavy atoms and applying an overlap threshold greater than -0.3 Å to define significant VDW interactions.

### Statistical analysis

All data represent pooled results from at least two, three, or five independent experiments, as indicated in the figure legends. Statistical analyses were performed using GraphPad Prism v10.0 (GraphPad Software). Normality and variance were assessed for each dataset to guide the choice of statistical tests, and two-sided tests were used unless otherwise specified. For time-course experiments, data were analyzed using two-way repeated-measures ANOVA or mixed-effects models, with time and treatment as factors and individual donors or experiments treated as repeated measures. Data were also analyzed using three-way repeated-measures ANOVA or mixed-effects models, with genotype, treatment, and time included as factors and individual donors or experiments treated as repeated measures. When appropriate, post hoc multiple-comparisons tests were performed following ANOVA. A p value < 0.05 was considered statistically significant for all analyses.

For circadian datasets, cosinor analysis ( COSINOR online v2.4) (53) was used to determine rhythm parameters (MESOR, amplitude, acrophase). Statistical significance was defined as p < 0.05 (*), p < 0.01 (**), p < 0.001 (***), and p < 0.0001 (****); ns, not significant.

Time-series data were analyzed using R (version 4.5.1). Circadian rhythmicity was modeled using the cosinor method, a regression-based technique for estimating rhythmic parameters (54) using the GLMM cosinor package (55). This approach allows for flexible modeling of rhythmic components across donors and conditions. A model was fitted to estimate a mesor, acrophase and amplitude value for each sample (a sample here is a unique combination of donor and treatment) with a period of 24 hour.

## Supplementary Figures

**Supplementary Figure 1. Related to Figure 1** (A) Representative flow cytometry plots and gating strategy of hMDM. Macrophages were defined as HLA-DR⁺ CD64⁺ CD14⁺ CD206⁺ CD163⁺ and negative for CD1a, CD3, CD15, CD19, and CD56 (3 donors were analyzed separately). (B) Experimental workflow for circadian analysis in human macrophages. PBMCs from five donors (D1-D5) were isolated, monocytes differentiated into macrophages with M-CSF (50 ng/mL, 7 d), and cells synchronized by a 2-h serum shock at phase-shifted starting times, enabling simultaneous collection of all circadian time points (CT12-CT48). Cells were primed with LPS (0.5 µg/mL, 4 h) and activated with nigericin (20 µM, 30-120 min).

**Supplementary Figure 2. Related to Figure 2** (A-B) HEK293T cells were transfected with FLAG-NLRP3 and CRY1 (A) or CRY2 (B) for 48 h. Anti-FLAG immunoprecipitation followed by immunoblotting was performed to detect CRY1 or CRY2 binding; actin served as a loading control (n=3). (C, E, F, H) Representative confocal images of PLA (red) in THP-1 NLRP3-KO (C), U937 WT (E) and NLRP3-KO (F), and PLA controls in hMDMs using single antibodies to assess nonspecific signal (H). Nuclei were stained with Hoechst. Images were acquired on an Opera Phenix HCS microscope (60× water objective); scale bar, 50 µm (n=2). (D, G, J) Immunoblots of NLRP3, CRY1, and CRY2 in THP-1 WT/NLRP3-KO (D), U937 WT/NLRP3-KO (G), and hMDMs (J). Actin served as loading control (n>5). (I) Quantification of PLA signal in hMDMs from (H). Single-antibody controls were compared with NLRP3-CRY1 or NLRP3-CRY2 pairs. ≥500 cells/condition; mean ± SD; Mann-Whitney, ****p < 0.0001. (K-L) HEK293T cells were transfected with CRY1-HA (J) or CRY2-HA (K) and the indicated FLAG constructs for 48 h prior to anti-FLAG IP and immunoblotting. Actin served as loading control (n=3). * indicates non-specific band.

**Supplementary Figure 3. Related to Figure 2** (A, C, E, G, I) Quantification of CRY2-NLRP3 PLA signal across circadian times in hMDMs from five donors under untreated conditions. Each color represents a different donor. (B, D, F, H, J) Corresponding individual cosinor fits for CRY2-NLRP3 PLA signal across circadian time for each donor. (L) Summary table of Kruskal-Wallis statistical analyses corresponding to panels (B-K). (M) Quantification of ATM-ATM PLA signal in samples from the same donor shown in Figure 2E. The table reports Kruskal-Wallis test results, indicating no significant variation across circadian time.

**Supplementary Figure 4. Related to Figure 3** (A) Immunoblot analysis of NLRP3, CRY1, CRY2, and GSDMD in THP-1 cells (n=3) treated with LPS (0.5 µg/mL, 4 h) with or without nigericin (10 µM, 1 h). Actin served as a loading control. (B) Representative PLA images (red) showing CRY1-NLRP3 or CRY2-NLRP3 interactions in THP-1 WT cells (N=3) primed with LPS (0.5 µg/mL, 4 h) and untreated or activated with nigericin (20 µM, 1 h). Nuclei were stained with Hoechst; images were acquired on an Opera Phenix HCS microscope using a 40× objective; scale bar, 50 µm. (C-D) Quantification of PLA total spot area per cell for CRY1-NLRP3 (C) and CRY2-NLRP3 (D). ≥500 cells per condition; mean ± SEM; Mann-Whitney test, ****p < 0.0001. (E-F) HEK293T cells were transfected with FLAG-NLRP3 and CRY1 (A) or CRY2 (B) for 48 h, treated with nigericin (10 µM, 1 h) or vehicle, and subjected to anti-FLAG immunoprecipitation followed by immunoblotting. Actin served as a loading control (n=4).

**Supplementary Figure 5. Related to Figure 3** (A, C, E, G, I) Quantification of CRY2-NLRP3 PLA signal across circadian times in hMDMs from five donors, comparing untreated conditions (data from Supplementary Figure 3) with nigericin-treated cells from the same donors. Each color represents a different donor. (B, D, F, H, J) Corresponding individual cosinor fits for CRY2-NLRP3 PLA signal across circadian time for each donor under untreated and nigericin-treated conditions. (K, L) Lollipop plots showing amplitude (K) and acrophase (L) of CRY2-NLRP3 interactions for each donor. Purple indicates untreated cells; pink indicates nigericin-treated cells.

**Supplementary Figure 6. Related to Figure 4** (A-M) hMDMs were differentiated for 7 days with M-CSF (50 ng/mL) and pretreated for 48 h with DMSO or CRY stabilizers KL001 (5 µM) (A,F, K, L), KL044 (5 µM) (B, F, M), or TH301 (2 µM) (C-E; G-J). Cells were then primed with LPS (0.5 µg/mL, 4 h) and activated with nigericin (20 µM, 1-1.5 h) or vehicle as indicated. (A-C) Protein expression of CRY1, CRY2, and NLRP3 was assessed by immunoblotting; actin served as a loading control (n=3). (D) representative time course of nigericin-induced cell death measured by DRAQ7 incorporation (Opera Phenix). Top: kinetic curves of DRAQ7 signal over time. Bottom left: quantification of cell death as area under the curve (AUC) (GraphPad Prism), shown as mean ± SD. Bottom right: paired individual values from three independent experiments, each represented by four pooled technical replicates (12 total points), Mann-Whitney, **p < 0.005. (E) Caspase-1 activity normalized to CV staining. Top panel shows data as mean ± SD from four independent experiments, each performed with three technical replicates (12 wells per condition). Bottom panel shows paired individual values from the four independent experiments. Statistical comparison was performed using a paired Mann-Whitney test (****p < 0.0001). (F-G) Representative confocal images of ASC specks (green) in hMDMs primed with LPS (0.5 µg/mL, 4 h) and left untreated (F) or activated with nigericin (20 µM, 1.5 h) (G). Nuclei were stained with Hoechst; images were acquired on an Opera Phenix HCS microscope using a 20× objective; scale bar, 50 µm; arrows indicate ASC specks. (H) Quantification of ASC-speck-positive cells from (G), shown as mean ± SD from three independent experiments, each performed with four experimental replicates (12 wells per condition), with ≥200 cells analyzed per condition. Statistical comparisons were performed using a paired Mann-Whitney test (*p = 0.0225, ****p < 0.0001). (I) hMDMs were primed with LPS (0.5 µg/mL, 4 h) and treated with nigericin (20 µM, 3 h). Levels of caspase-1 and IL-1β were analyzed by immunoblotting in cell lysates and corresponding supernatants (n = 2). (J) IL-1β secretion measured by ELISA in supernatants from hMDMs treated with nigericin (20 µM, 3 h). Top: pooled quantification from six measurements obtained across three independent experiments, shown as mean ± SD. Bottom: paired individual values from the three independent experiments. Statistical comparisons were performed using a unpaired Mann-Whitney test. (K) Representative PLA images (red) showing CRY1-NLRP3 interactions in hMDMs pretreated with DMSO, KL001 (5 µM) or KL044 (5 µM), primed with LPS (0.5 µg/mL, 4 h) and left untreated or activated with nigericin (20 µM, 1 h). Nuclei were stained with Hoechst; images were acquired on an Opera Phenix HCS microscope using a 40× objective; scale bar, 50 µm. (L-M) Quantification of PLA signal for CRY1-NLRP3 (total spot area per cell, Harmony software). Left: representative experiment showing three wells per condition with ≥1,000 cells analyzed per well. Right: paired summary of three independent experiments (n = 3). Data are shown as paired dots with mean ± SD. Statistical significance was determined using a paired Mann-Whitney test (****p < 0.0001). (N-P) hMDMs were primed with LPS (0.5 µg/mL, 4 h), pretreated with MCC950 (150 nM, 30 min) before nigericin treatment (20 µM). (N) Cell death was monitored by DRAQ7 incorporation using an Opera Phenix HCS microscope (objective x20 air). (O) The area under the curve (AUC) for cell death was quantified across circadian times. (L) Caspase-1 activity was quantified and normalized to Crystal Violet staining (n=3).

**Supplementary Figure 7. Related to Figure 6** (A) Schematic representation of FLAG-NLRP3 full-length (FL) and truncated constructs containing the PYD, NACHT, or LRR domains, or PYD+NACHT (NLRP3-short). (B) Schematic representation of FLAG-CRY2 full-length (FL) and truncated constructs containing the PHR or C-terminal (CT) domains. (C) HEK293T cells were transfected with FLAG-NLRP3 constructs and CRY1 for 48 h before anti-FLAG immunoprecipitation and immunoblotting (n=3). (D) Structural alignments of AlphaFold-predicted models aligned with published structures. Left panel: human NLRP3 (AlphaFold) aligned with the experimental human NLRP3 structure (PDB 7PZC). Middle panel: human CRY2 (AlphaFold) aligned with mouse CRY2 (PDB 4I6J). Right panel: AlphaFold-predicted CRY2-NLRP3 complex aligned with human NLRP3 (PDB 7PZC) and mouse CRY2 (PDB 4I6J). The NACHT and LRR domains of NLRP3 are shown in different shades of blue, the PYD domain in gray, mouse CRY2 in red, the AlphaFold NLRP3 model in dark blue, and the AlphaFold CRY2 model in orange. (E) Left panel: AlphaFold-predicted model of the human CRY1-NLRP3 complex. Right panel: structural alignment of AlphaFold-predicted CRY1 with the human NLRP3 structure (PDB 7PZC). NLRP3 domains are color-coded as follows: PYD (gray), NACHT (dark blue), and LRR (light blue); CRY1 structure is shown in purple. (F-G) HEK293T cells were transfected with FLAG-CRY1 and WT NLRP3 or CAPS variants (V262G, R260W, D303N, M701T) for 48 h, followed by anti-FLAG immunoprecipitation and immunoblotting (n=3). * indicates non-specific band.

**Supplementary Figure 8. Related to Figure 7** (A) hMDMs were synchronized as in Figure 5, primed with LPS (0.5 µg/mL, 4 h), and pretreated or not with MCC950 (150 nM, 30 min) before stimulation with nigericin (20 µM). Cell death was monitored in real time by DRAQ7 incorporation using an Opera Phenix high-content imaging system. (B) Q-Q plot assessing normality of the IL-1β data from (Fig 7F-I). (C) Results of Sidak’s post hoc multiple comparisons following ANOVA in Fig7J. (D) Caspase-1 activity normalized to crystal violet staining in synchronized U937 cells expressing NLRP3 WT, M701T or Q703K. (E-F) Caspase-1 activity in synchronized U937 cells expressing NLRP3 WT (E) or the V262G mutant (F), measured at 24, 30, 36, and 42 h post-synchronization and treated or not with MCC950. Top: caspase-1 activity across time. Bottom: corresponding two-way ANOVA summary tables. (G) Q-Q plots assessing normality of residuals for the two-way ANOVA analyses shown in (E) and (F) with results from Sidak’s post hoc multiple comparisons (H). (I-K) Caspase-1 activity in synchronized U937 cells expressing NLRP3 WT (I), M701T (J) or Q703K (K) displayed as in (E-F), with corresponding two-way ANOVA summary table shown below (M). (L) Q-Q plots assessing normality of residuals for the two-way ANOVA analyses shown in (I-K). (M) Summary table reporting three-way ANOVA results for caspase-1 activity across all U937 genotypes, with time, MCC950 treatment, and genotype as factors. (N) Sidak’s multiple comparisons results calculated from the three-way ANOVA in (M).

## Acknowledgments

We are grateful to the staff of the Cell Imaging Platform of the CRCL (PIC, CRCL, Lyon) and especially Christophe Vanbelle for their training, assistance and support. We also thank Stéphane Giraud from the Center for drug discovery and development (C3D) Platform of the CRCL for insightful discussions and guidance regarding AlphaFold structural analyses. We are also grateful to Kiran Padmanabhan for critical reading of the manuscript and constructive feedbacks.

## Bibliography

1. Scheiermann C, Gibbs J, Ince L, Loudon A. Clocking in to immunity. Nat Rev Immunol. juill 2018;18(7):423–37. doi:10.1038/s41577-018-0008-4 PubMed PMID: 29662121.

2. Holtkamp SJ, Ince LM, Barnoud C, Schmitt MT, Sinturel F, Pilorz V, et al. Circadian clocks guide dendritic cells into skin lymphatics. Nat Immunol. nov 2021;22(11):1375–81. doi:10.1038/s41590-021-01040-x PubMed PMID: 34663979; PubMed Central PMCID: PMC8553624.

3. Silver AC, Arjona A, Walker WE, Fikrig E. The circadian clock controls toll-like receptor 9-mediated innate and adaptive immunity. Immunity. 24 févr 2012;36(2):251–61. doi:10.1016/j.immuni.2011.12.017 PubMed PMID: 22342842; PubMed Central PMCID: PMC3315694.

4. Curtis AM, Bellet MM, Sassone-Corsi P, O’Neill LAJ. Circadian clock proteins and immunity. Immunity. 20 févr 2014;40(2):178–86. doi:10.1016/j.immuni.2014.02.002 PubMed PMID: 24560196.

5. Keller M, Mazuch J, Abraham U, Eom GD, Herzog ED, Volk HD, et al. A circadian clock in macrophages controls inflammatory immune responses. Proc Natl Acad Sci U S A. 15 déc 2009;106(50):21407–12. doi:10.1073/pnas.0906361106 PubMed PMID: 19955445; PubMed Central PMCID: PMC2795539.

6. Labrecque N, Cermakian N. Circadian Clocks in the Immune System. J Biol Rhythms. août 2015;30(4):277–90. doi:10.1177/0748730415577723 PubMed PMID: 25900041.

7. Hand LE, Hopwood TW, Dickson SH, Walker AL, Loudon ASI, Ray DW, et al. The circadian clock regulates inflammatory arthritis. FASEB J Off Publ Fed Am Soc Exp Biol. nov 2016;30(11):3759–70. doi:10.1096/fj.201600353R PubMed PMID: 27488122; PubMed Central PMCID: PMC5067252.

8. Pariollaud M, Gibbs JE, Hopwood TW, Brown S, Begley N, Vonslow R, et al. Circadian clock component REV-ERBα controls homeostatic regulation of pulmonary inflammation. J Clin Invest. 1 juin 2018;128(6):2281-96. doi:10.1172/JCI93910 PubMed PMID: 29533925; PubMed Central PMCID: PMC5983347.

9. Partch CL, Green CB, Takahashi JS. Molecular architecture of the mammalian circadian clock. Trends Cell Biol. févr 2014;24(2):90–9. doi:10.1016/j.tcb.2013.07.002 PubMed PMID: 23916625; PubMed Central PMCID: PMC3946763.

10. Liu Y, Sancar A. Biochemical mechanism of the mammalian circadian clock. FEBS Lett. 25 août 2025. doi:10.1002/1873-3468.70150 PubMed PMID: 40854106; PubMed Central PMCID: PMC12380416.

11. Ruben MD, Wu G, Smith DF, Schmidt RE, Francey LJ, Lee YY, et al. A database of tissue-specific rhythmically expressed human genes has potential applications in circadian medicine. Sci Transl Med. 12 sept 2018;10(458):eaat8806. doi:10.1126/scitranslmed.aat8806 PubMed PMID: 30209245; PubMed Central PMCID: PMC8961342.

12. Ikeda R, Tsuchiya Y, Koike N, Umemura Y, Inokawa H, Ono R, et al. REV-ERBα and REV-ERBβ function as key factors regulating Mammalian Circadian Output. Sci Rep. 15 juill 2019;9(1):10171. doi:10.1038/s41598-019-46656-0 PubMed PMID: 31308426; PubMed Central PMCID: PMC6629614.

13. Mure LS, Le HD, Benegiamo G, Chang MW, Rios L, Jillani N, et al. Diurnal transcriptome atlas of a primate across major neural and peripheral tissues. Science. 16 mars 2018;359(6381):eaao0318. doi:10.1126/science.aao0318 PubMed PMID: 29439024; PubMed Central PMCID: PMC5924732.

14. Narasimamurthy R, Hatori M, Nayak SK, Liu F, Panda S, Verma IM. Circadian clock protein cryptochrome regulates the expression of proinflammatory cytokines. Proc Natl Acad Sci U S A. 31 juill 2012;109(31):12662–7. doi:10.1073/pnas.1209965109 PubMed PMID: 22778400; PubMed Central PMCID: PMC3411996.

15. Hashiramoto A, Yamane T, Tsumiyama K, Yoshida K, Komai K, Yamada H, et al. Mammalian clock gene Cryptochrome regulates arthritis via proinflammatory cytokine TNF-alpha. J Immunol Baltim Md 1950. 1 févr 2010;184(3):1560–5. doi:10.4049/jimmunol.0903284 PubMed PMID: 20042581.

16. Kondratov RV, Kondratova AA, Gorbacheva VY, Vykhovanets OV, Antoch MP. Early aging and age-related pathologies in mice deficient in BMAL1, the core componentof the circadian clock. Genes Dev. 15 juill 2006;20(14):1868–73. doi:10.1101/gad.1432206 PubMed PMID: 16847346; PubMed Central PMCID: PMC1522083.

17. Lamia KA, Sachdeva UM, DiTacchio L, Williams EC, Alvarez JG, Egan DF, et al. AMPK regulates the circadian clock by cryptochrome phosphorylation and degradation. Science. 16 oct 2009;326(5951):437–40. doi:10.1126/science.1172156 PubMed PMID: 19833968; PubMed Central PMCID: PMC2819106.

18. Yoo SH, Mohawk JA, Siepka SM, Shan Y, Huh SK, Hong HK, et al. Competing E3 ubiquitin ligases govern circadian periodicity by degradation of CRY in nucleus and cytoplasm. Cell. 28 févr 2013;152(5):1091–105. doi:10.1016/j.cell.2013.01.055 PubMed PMID: 23452855; PubMed Central PMCID: PMC3694781.

19. Huber AL, Papp SJ, Chan AB, Henriksson E, Jordan SD, Kriebs A, et al. CRY2 and FBXL3 Cooperatively Degrade c-MYC. Mol Cell. 17 nov 2016;64(4):774–89. doi:10.1016/j.molcel.2016.10.012 PubMed PMID: 27840026; PubMed Central PMCID: PMC5123859.

20. Martinon F, Burns K, Tschopp J. The inflammasome: a molecular platform triggering activation of inflammatory caspases and processing of proIL-beta. Mol Cell. août 2002;10(2):417–26. doi:10.1016/s1097-2765(02)00599-3 PubMed PMID: 12191486.

21. Pétrilli V, Dostert C, Muruve DA, Tschopp J. The inflammasome: a danger sensing complex triggering innate immunity. Curr Opin Immunol. déc 2007;19(6):615–22. doi:10.1016/j.coi.2007.09.002 PubMed PMID: 17977705.

22. Xiao L, Magupalli VG, Wu H. Cryo-EM structures of the active NLRP3 inflammasome disc. Nature. janv 2023;613(7944):595–600. doi:10.1038/s41586-022-05570-8 PubMed PMID: 36442502; PubMed Central PMCID: PMC10091861.

23. Hochheiser IV, Pilsl M, Hagelueken G, Moecking J, Marleaux M, Brinkschulte R, et al. Structure of the NLRP3 decamer bound to the cytokine release inhibitor CRID3. Nature. avr 2022;604(7904):184–9. doi:10.1038/s41586-022-04467-w PubMed PMID: 35114687.

24. Broz P, Dixit VM. Inflammasomes: mechanism of assembly, regulation and signalling. Nat Rev Immunol. juill 2016;16(7):407–20. doi:10.1038/nri.2016.58 PubMed PMID: 27291964.

25. Nadjar J, Monnier S, Bastien E, Huber AL, Oddou C, Bardoulet L, et al. Optogenetically controlled inflammasome activation demonstrates two phases of cell swelling during pyroptosis. Sci Signal. 23 avr 2024;17(833):eabn8003. doi:10.1126/scisignal.abn8003 PubMed PMID: 38652763.

26. Mangan MSJ, Olhava EJ, Roush WR, Seidel HM, Glick GD, Latz E. Targeting the NLRP3 inflammasome in inflammatory diseases. Nat Rev Drug Discov. août 2018;17(8):588–606. doi:10.1038/nrd.2018.97 PubMed PMID: 30026524.

27. Juliana C, Fernandes-Alnemri T, Kang S, Farias A, Qin F, Alnemri ES. Non-transcriptional priming and deubiquitination regulate NLRP3 inflammasome activation. J Biol Chem. 19 oct 2012;287(43):36617–22. doi:10.1074/jbc.M112.407130 PubMed PMID: 22948162; PubMed Central PMCID: PMC3476327.

28. Py BF, Kim MS, Vakifahmetoglu-Norberg H, Yuan J. Deubiquitination of NLRP3 by BRCC3 critically regulates inflammasome activity. Mol Cell. 24 janv 2013;49(2):331–8. doi:10.1016/j.molcel.2012.11.009 PubMed PMID: 23246432.

29. O’Keefe ME, Dubyak GR, Abbott DW. Post-translational control of NLRP3 inflammasome signaling. J Biol Chem. juin 2024;300(6):107386. doi:10.1016/j.jbc.2024.107386 PubMed PMID: 38763335; PubMed Central PMCID: PMC11245928.

30. Niu T, De Rosny C, Chautard S, Rey A, Patoli D, Groslambert M, et al. NLRP3 phosphorylation in its LRR domain critically regulates inflammasome assembly. Nat Commun. 6 oct 2021;12(1):5862. doi:10.1038/s41467-021-26142-w PubMed PMID: 34615873; PubMed Central PMCID: PMC8494922.

31. Paik S, Kim JK, Silwal P, Sasakawa C, Jo EK. An update on the regulatory mechanisms of NLRP3 inflammasome activation. Cell Mol Immunol. mai 2021;18(5):1141–60. doi:10.1038/s41423-021-00670-3 PubMed PMID: 33850310; PubMed Central PMCID: PMC8093260.

32. Early JO, Menon D, Wyse CA, Cervantes-Silva MP, Zaslona Z, Carroll RG, et al. Circadian clock protein BMAL1 regulates IL-1β in macrophages via NRF2. Proc Natl Acad Sci U S A. 4 sept 2018;115(36):E8460–8. doi:10.1073/pnas.1800431115 PubMed PMID: 30127006; PubMed Central PMCID: PMC6130388.

33. Pourcet B, Zecchin M, Ferri L, Beauchamp J, Sitaula S, Billon C, et al. Nuclear Receptor Subfamily 1 Group D Member 1 Regulates Circadian Activity of NLRP3 Inflammasome to Reduce the Severity of Fulminant Hepatitis in Mice. Gastroenterology. avr 2018;154(5):1449–1464.e20. doi:10.1053/j.gastro.2017.12.019 PubMed PMID: 29277561; PubMed Central PMCID: PMC5892845.

34. Wang S, Lin Y, Yuan X, Li F, Guo L, Wu B. REV-ERBα integrates colon clock with experimental colitis through regulation of NF-κB/NLRP3 axis. Nat Commun. 12 oct 2018;9(1):4246. doi:10.1038/s41467-018-06568-5 PubMed PMID: 30315268; PubMed Central PMCID: PMC6185905.

35. O’Siorain JR, Cox SL, Payet C, Nally FK, He Y, Drewinksi TT, et al. Time-of-day control of mitochondria regulates NLRP3 inflammasome activation in macrophages. FASEB J Off Publ Fed Am Soc Exp Biol. 13 déc 2024;38(24):e70235. doi:10.1096/fj.202400508RR PubMed PMID: 39686706; PubMed Central PMCID: PMC11669068.

36. Hirota T, Lee JW, St John PC, Sawa M, Iwaisako K, Noguchi T, et al. Identification of small molecule activators of cryptochrome. Science. 31 août 2012;337(6098):1094–7. doi:10.1126/science.1223710 PubMed PMID: 22798407; PubMed Central PMCID: PMC3589997.

37. Lee JW, Hirota T, Kumar A, Kim NJ, Irle S, Kay SA. Development of Small-Molecule Cryptochrome Stabilizer Derivatives as Modulators of the Circadian Clock. ChemMedChem. sept 2015;10(9):1489–97. doi:10.1002/cmdc.201500260 PubMed PMID: 26174033; PubMed Central PMCID: PMC4576822.

38. Miller S, Son YL, Aikawa Y, Makino E, Nagai Y, Srivastava A, et al. Isoform-selective regulation of mammalian cryptochromes. Nat Chem Biol. juin 2020;16(6):676–85. doi:10.1038/s41589-020-0505-1 PubMed PMID: 32231341.

39. Coll RC, Hill JR, Day CJ, Zamoshnikova A, Boucher D, Massey NL, et al. MCC950 directly targets the NLRP3 ATP-hydrolysis motif for inflammasome inhibition. Nat Chem Biol. juin 2019;15(6):556–9. doi:10.1038/s41589-019-0277-7 PubMed PMID: 31086327.

40. Xing W, Busino L, Hinds TR, Marionni ST, Saifee NH, Bush MF, et al. SCF(FBXL3) ubiquitin ligase targets cryptochromes at their cofactor pocket. Nature. 4 avr 2013;496(7443):64–8. doi:10.1038/nature11964 PubMed PMID: 23503662; PubMed Central PMCID: PMC3618506.

41. Cosson C, Riou R, Patoli D, Niu T, Rey A, Groslambert M, et al. Functional diversity of NLRP3 gain-of-function mutants associated with CAPS autoinflammation. J Exp Med. 6 mai 2024;221(5):e20231200. doi:10.1084/jem.20231200 PubMed PMID: 38530241; PubMed Central PMCID: PMC10966137.

42. Hoffman HM, Mueller JL, Broide DH, Wanderer AA, Kolodner RD. Mutation of a new gene encoding a putative pyrin-like protein causes familial cold autoinflammatory syndrome and Muckle-Wells syndrome. Nat Genet. nov 2001;29(3):301–5. doi:10.1038/ng756 PubMed PMID: 11687797; PubMed Central PMCID: PMC4322000.

43. Putnam CD, Broderick L, Hoffman HM. The discovery of NLRP3 and its function in cryopyrin-associated periodic syndromes and innate immunity. Immunol Rev. mars 2024;322(1):259–82. doi:10.1111/imr.13292 PubMed PMID: 38146057; PubMed Central PMCID: PMC10950545.

44. Coll RC, Schroder K, Pelegrín P. NLRP3 and pyroptosis blockers for treating inflammatory diseases. Trends Pharmacol Sci. août 2022;43(8):653–68. doi:10.1016/j.tips.2022.04.003 PubMed PMID: 35513901.

45. Tsuji G, Hashimoto-Hachiya A, Yen VH, Takemura M, Yumine A, Furue K, et al. Metformin inhibits IL-1β secretion via impairment of NLRP3 inflammasome in keratinocytes: implications for preventing the development of psoriasis. Cell Death Discov. 2020;6:11. doi:10.1038/s41420-020-0245-8 PubMed PMID: 32194991; PubMed Central PMCID: PMC7055596.

46. Xian H, Liu Y, Rundberg Nilsson A, Gatchalian R, Crother TR, Tourtellotte WG, et al. Metformin inhibition of mitochondrial ATP and DNA synthesis abrogates NLRP3 inflammasome activation and pulmonary inflammation. Immunity. 13 juill 2021;54(7):1463–1477.e11. doi:10.1016/j.immuni.2021.05.004 PubMed PMID: 34115964; PubMed Central PMCID: PMC8189765.

47. Wagner PM, Fornasier SJ, Guido ME. Pharmacological Modulation of the Cytosolic Oscillator Affects Glioblastoma Cell Biology. Cell Mol Neurobiol. 22 juin 2024;44(1):51. doi:10.1007/s10571-024-01485-2 PubMed PMID: 38907776; PubMed Central PMCID: PMC11193694.

48. Chan P, Nagai Y, Wu Q, Hovsepyan A, Mkhitaryan S, Wang J, et al. Advancing Clinical Response Against Glioblastoma: Evaluating SHP1705 CRY2 Activator Efficacy in Preclinical Models and Safety in Phase I Trials. BioRxiv Prepr Serv Biol. 21 sept 2024;2024.09.17.613520. doi:10.1101/2024.09.17.613520 PubMed PMID: 39345648; PubMed Central PMCID: PMC11429762.

49. Abramson J, Adler J, Dunger J, Evans R, Green T, Pritzel A, et al. Accurate structure prediction of biomolecular interactions with AlphaFold 3. Nature. 2024;630(8016):493–500. doi:10.1038/s41586-024-07487-w PubMed PMID: 38718835; PubMed Central PMCID: PMC11168924.

50. Pettersen EF, Goddard TD, Huang CC, Meng EC, Couch GS, Croll TI, et al. UCSF ChimeraX: Structure visualization for researchers, educators, and developers. Protein Sci Publ Protein Soc. janv 2021;30(1):70–82. doi:10.1002/pro.3943 PubMed PMID: 32881101; PubMed Central PMCID: PMC7737788.

51. Goddard TD, Huang CC, Meng EC, Pettersen EF, Couch GS, Morris JH, et al. UCSF ChimeraX: Meeting modern challenges in visualization and analysis. Protein Sci Publ Protein Soc. janv 2018;27(1):14–25. doi:10.1002/pro.3235 PubMed PMID: 28710774; PubMed Central PMCID: PMC5734306.

52. Meng EC, Goddard TD, Pettersen EF, Couch GS, Pearson ZJ, Morris JH, et al. UCSF ChimeraX: Tools for structure building and analysis. Protein Sci Publ Protein Soc. nov 2023;32(11):e4792. doi:10.1002/pro.4792 PubMed PMID: 37774136; PubMed Central PMCID: PMC10588335.

53. Molcan L. Time distributed data analysis by Cosinor.Online application. BioRxiv. 12 déc 2016:597. 10.1101/805960

54. Cornelissen G. Cosinor-based rhythmometry. Theor Biol Med Model. 11 avr 2014;11:16. doi:10.1186/1742-4682-11-16 PubMed PMID: 24725531; PubMed Central PMCID: PMC3991883.

55. Parsons R, Jayasinghe O, White N, Chunduri P, Rawashdeh O. GLMMcosinor: Flexible Cosinor Modeling to Characterize Rhythmic Time Series Using a Generalized Linear Mixed Modeling Framework [Internet]. Bioinformatics; 2024 [cité 7 oct 2025]. Disponible sur: http://biorxiv.org/lookup/doi/10.1101/2024.04.10.588934 doi:10.1101/2024.04.10.588934

