## Supplementary figures and images for "CRY-NLRP3 complexes define a circadian checkpoint controlling inflammasome activation"

### Supplementary Figure 1

#### Figure S1

*Bardoulet et al.*

**A**

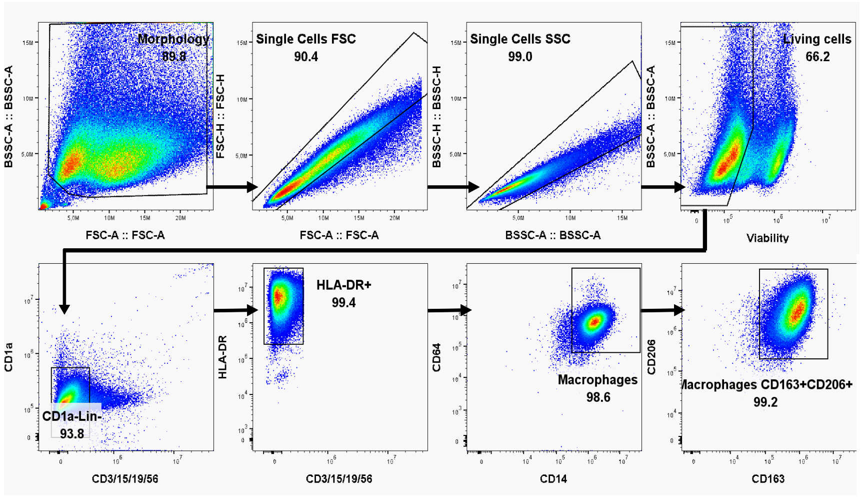


**B**

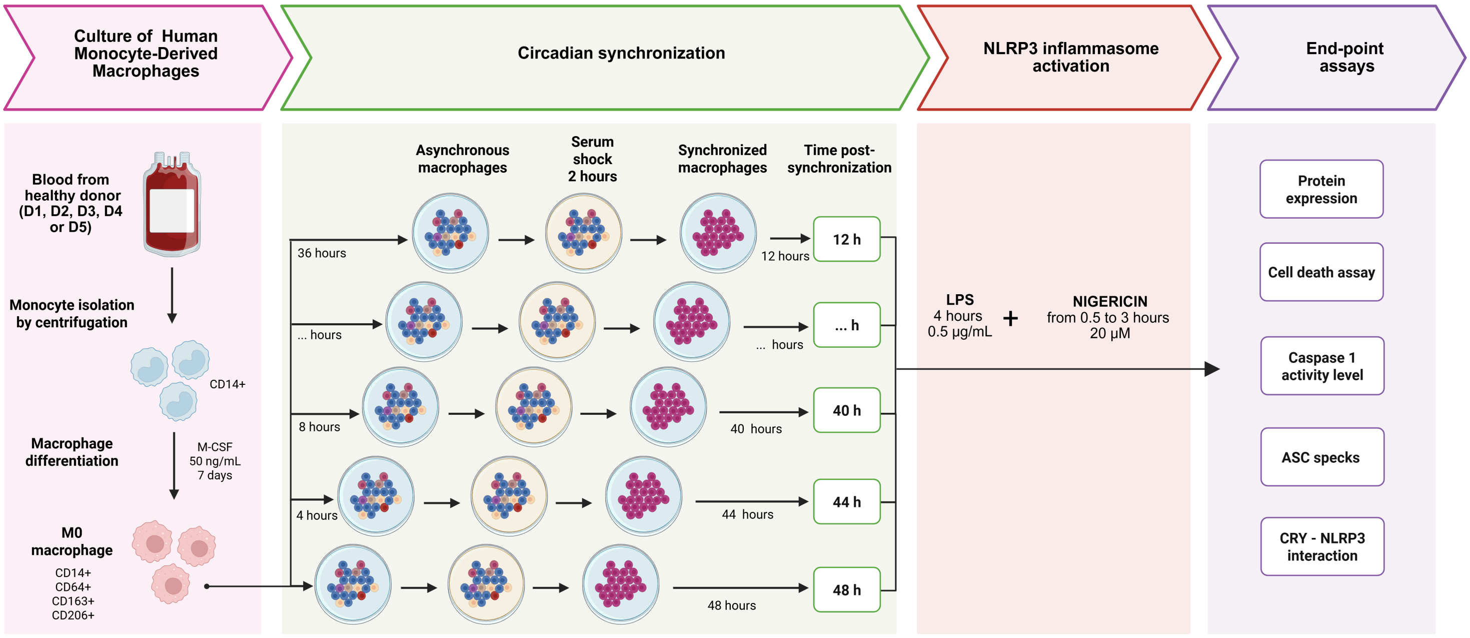

### Supplementary Figure 2

#### Figure S2

Bardoulet et al.

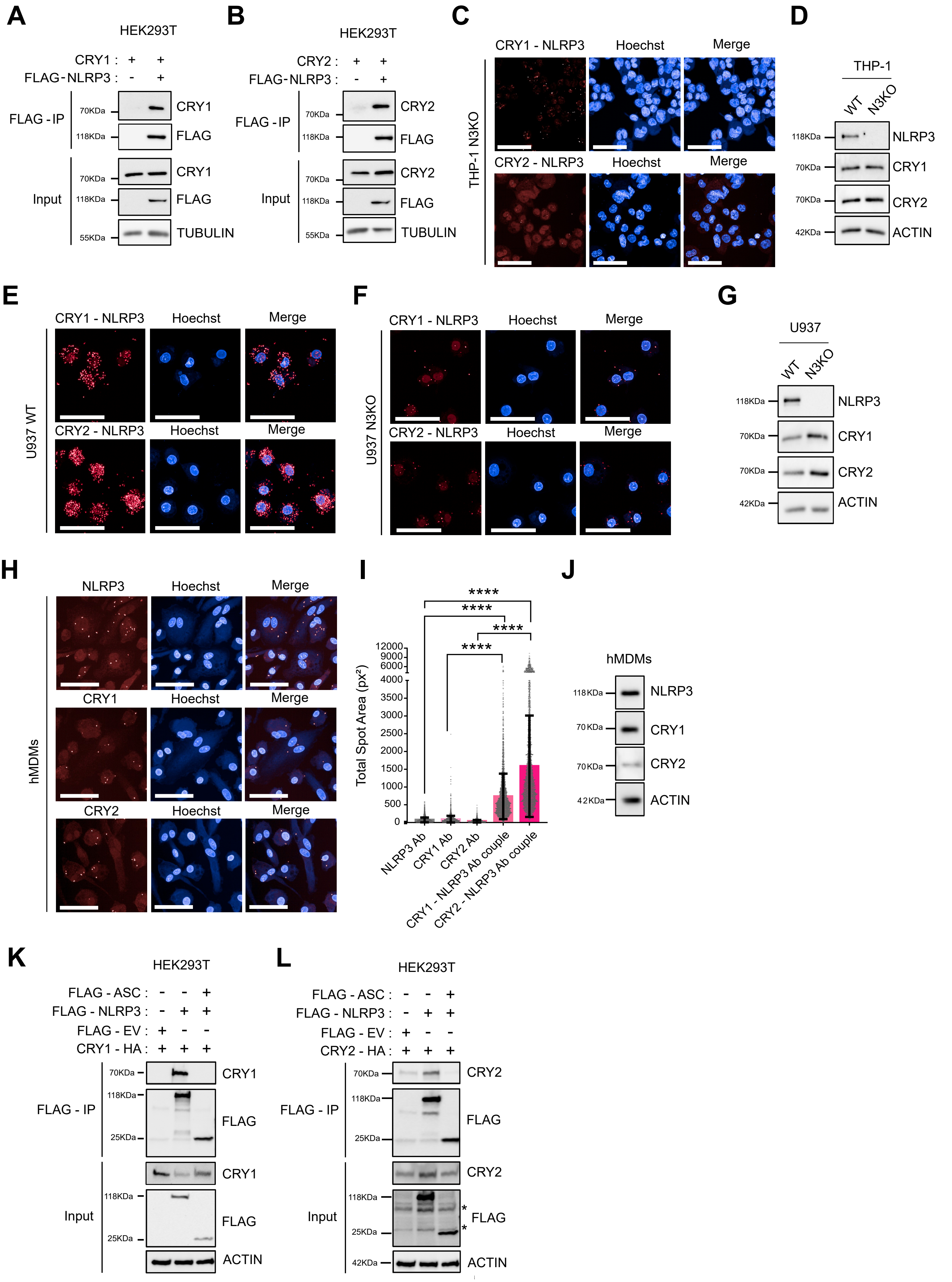

### Supplementary Figure 4

#### Figure S4

*Bardoulet et al.*

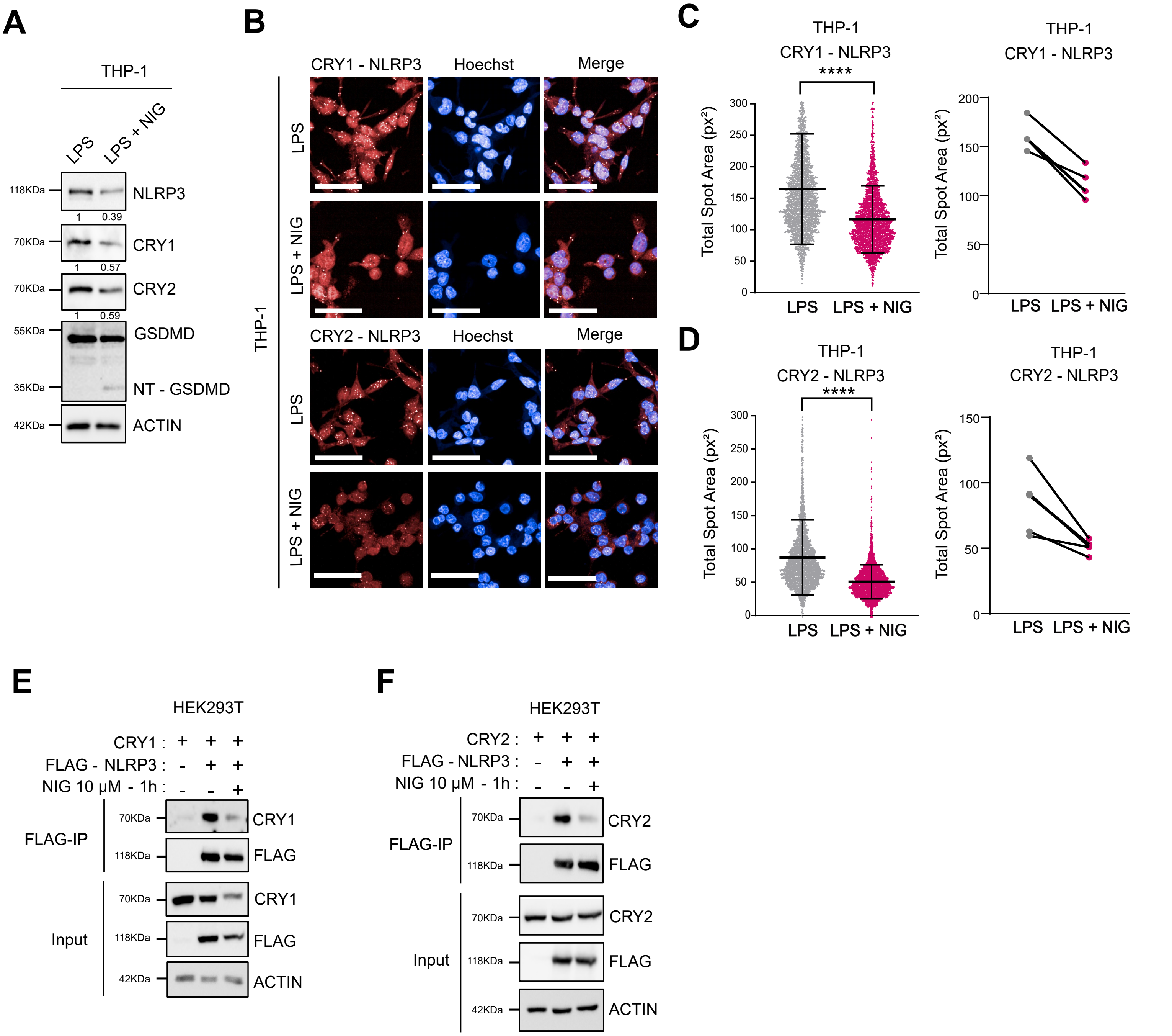

### Supplementary Figure 5

Figure S5

*Bardoulet et al.*

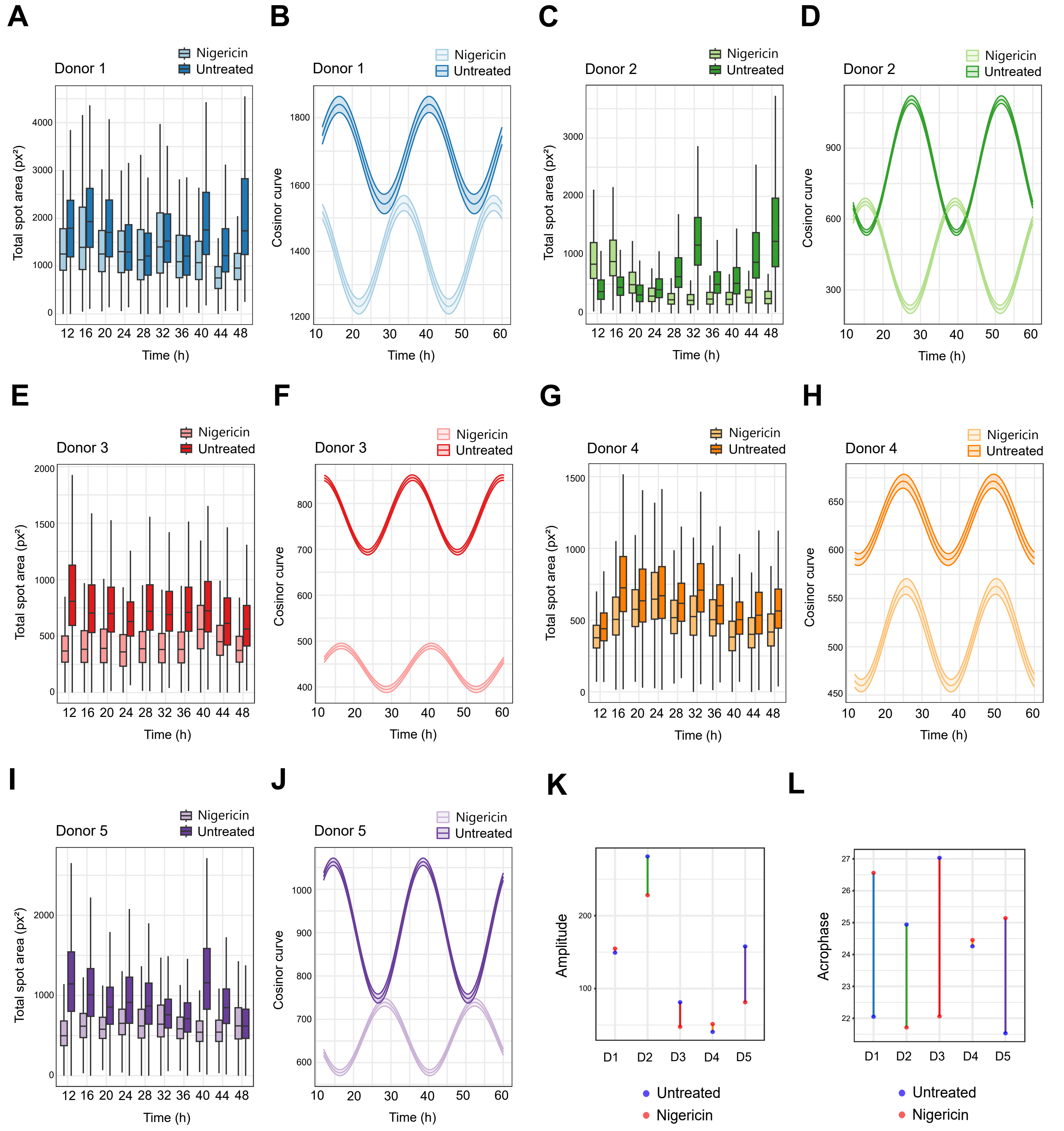

### Supplementary Figure 6

Figure S6

Bardoulet et al.

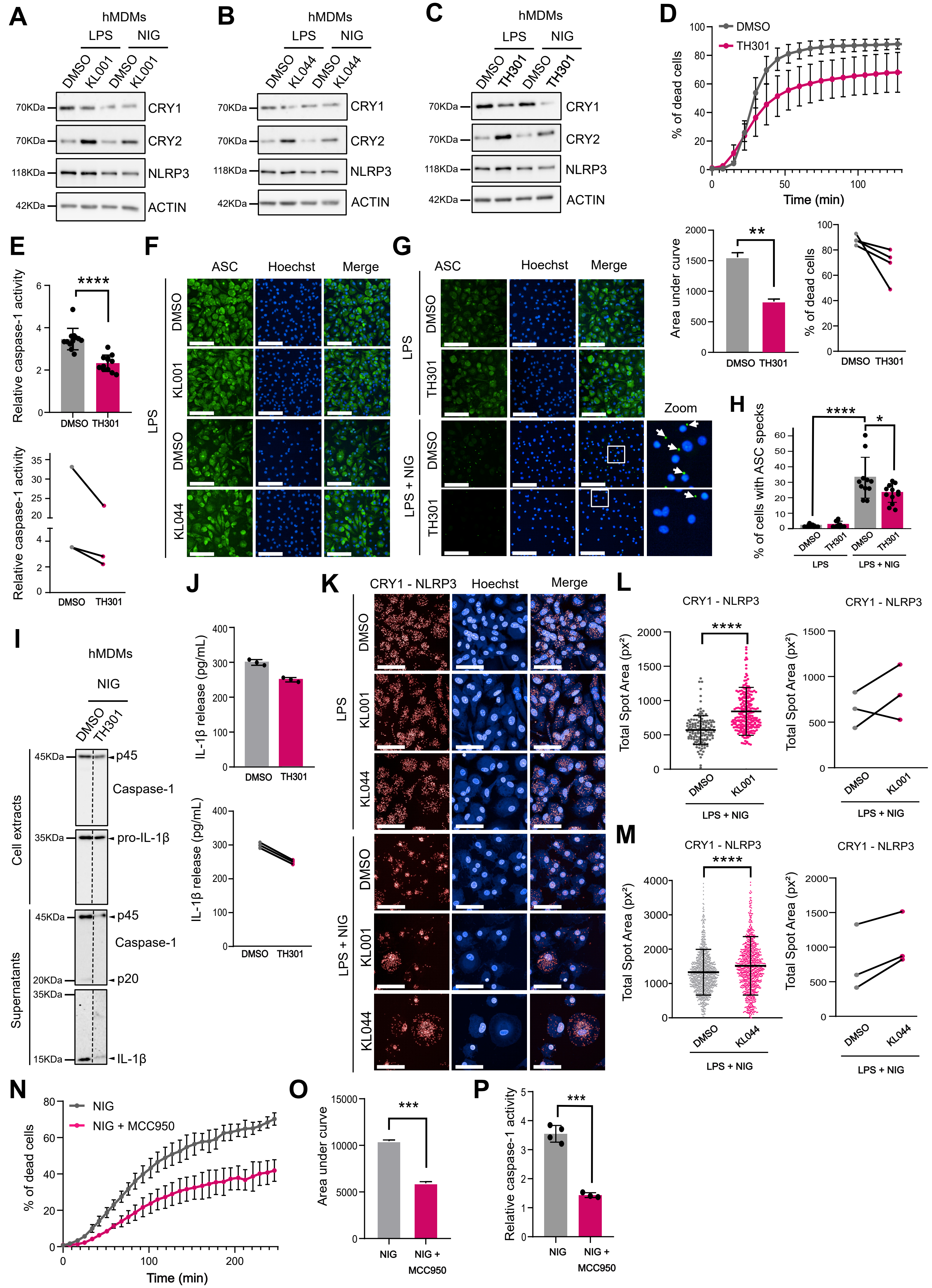

### Supplementary Figure 7

Figure S7

*Bardoulet et al.*

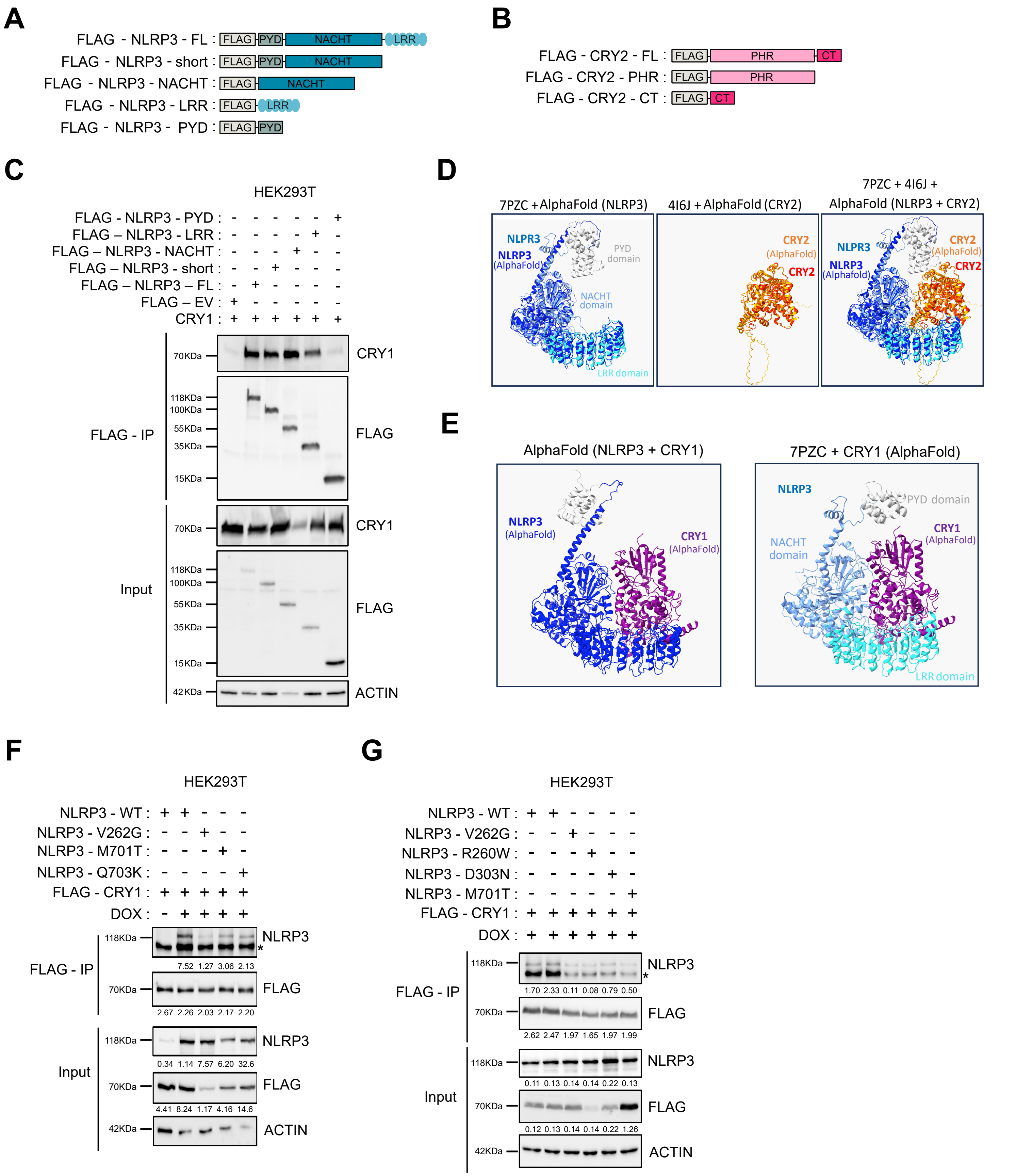
