## Supplementary Figure 3 for "CRY-NLRP3 complexes define a circadian checkpoint controlling inflammasome activation"

### Figure S3

*Bardoulet et al.*

A B C D

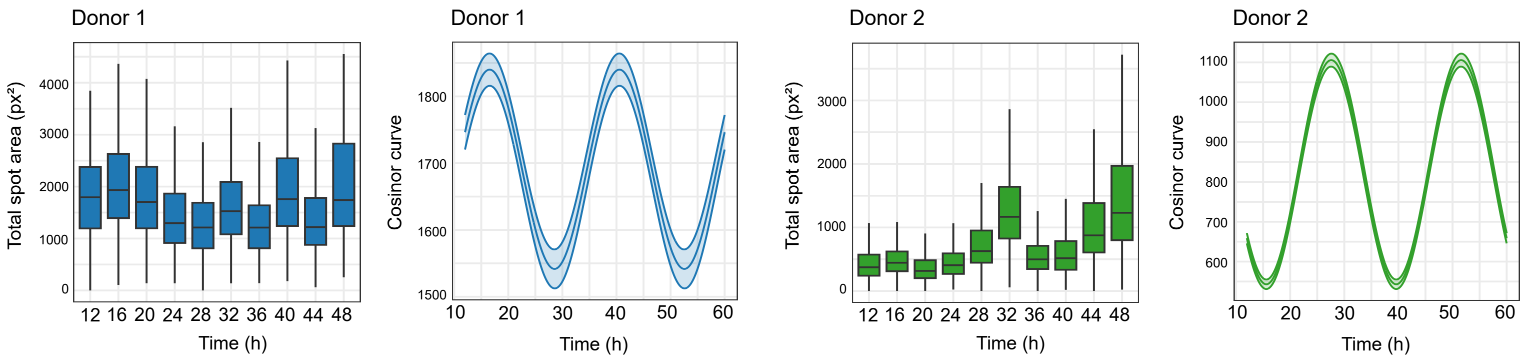


E F G H

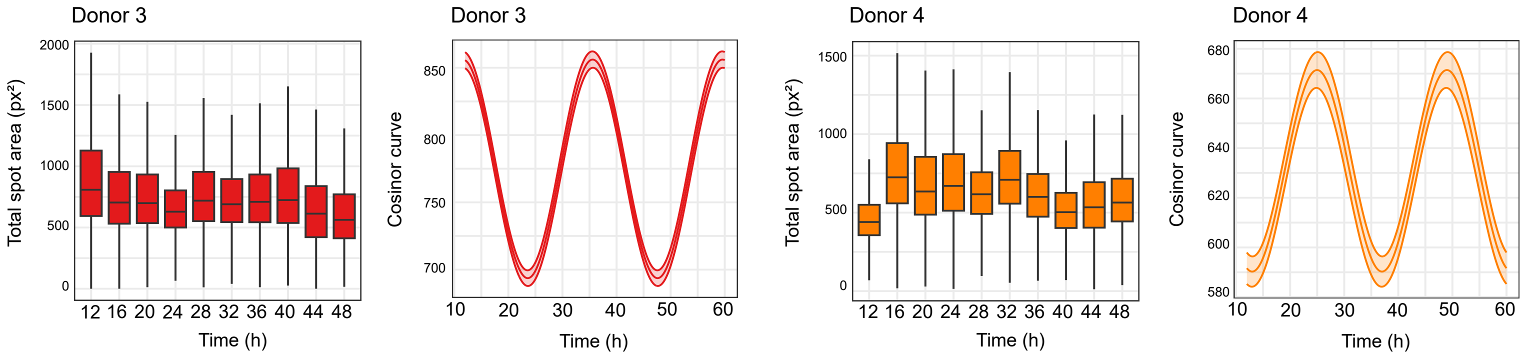


I J K

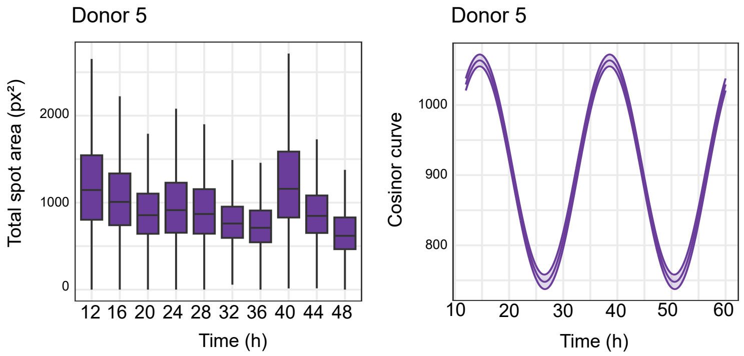


| Donneur | Kruskal-Wallis chi-squared | df | p-value |
|---|---|---|---|
| D1 | 276.7095 | 9 | $2.25 \times 10^{-54}$ |
| D2 | 3503.6085 | 9 | $p < 2.2 \times 10^{-16}$ |
| D3 | 2213.9088 | 9 | $p < 2.2 \times 10^{-16}$ |
| D4 | 5967.0393 | 9 | $p < 2.2 \times 10^{-16}$ |
| D5 | 3580.7445 | 9 | $p < 2.2 \times 10^{-16}$ |

L

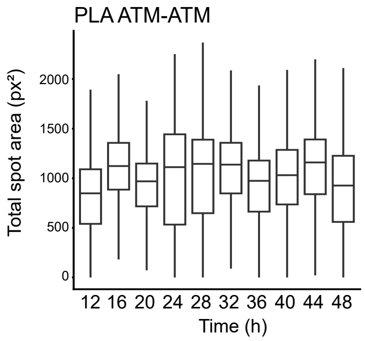


| | p-value | Significance |
|---|---|---|
| Amplitude | 0.408 | NS |
| Acrophase | 0.451 | NS |
