## Supplementary Figure 8 for "CRY-NLRP3 complexes define a circadian checkpoint controlling inflammasome activation"

### Figure S8

Bardoulet et al.

**A**

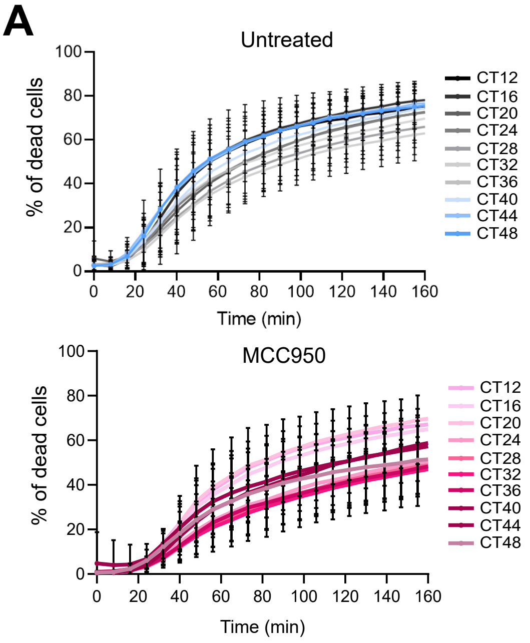


**B**

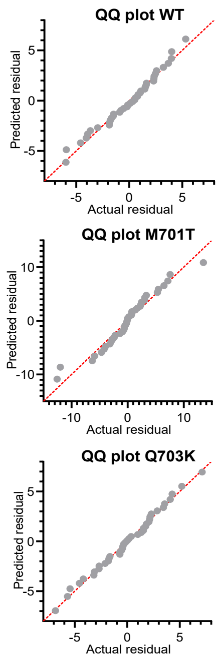


**C**

| | WT | | M701T | | Q703K | |
|---|---|---|---|---|---|---|
| Time (h) | p-value | p-value summary | p-value | p-value summary | p-value | p-value summary |
| 24 | 0,0334 | * | 0,0744 | ns | 0,2663 | ns |
| 30 | 0,9994 | ns | 0,0205 | * | 0,0504 | ns |
| 36 | 0,5949 | ns | 0,0129 | * | 0,0006 | *** |
| 42 | 0,248 | ns | 0,0024 | ** | 0,0543 | ns |
| 48 | 0,0878 | ns | 0,0509 | ns | 0,9186 | ns |

**D**

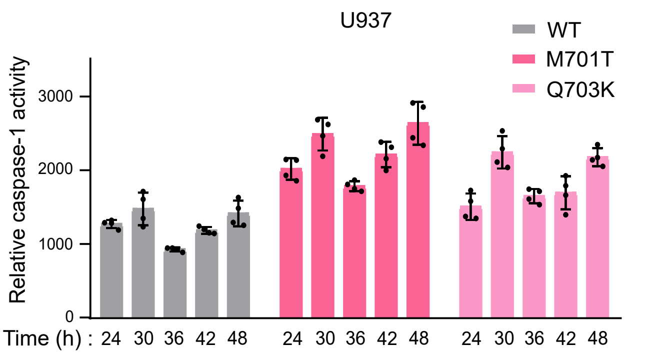


**E**

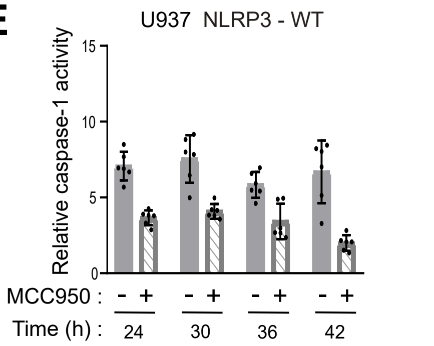


| 2-way ANOVA | p-value | p-value summary |
|---|---|---|
| Time | 0,0053 | ** |
| MCC950 | <0,0001 | **** |
| Time x MCC950 | 0,0347 | * |

**F**

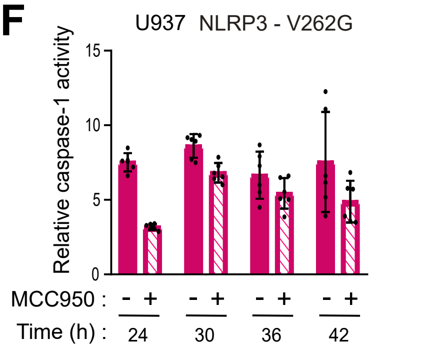


| 2-way ANOVA | p-value | p-value summary |
|---|---|---|
| Time | 0,0141 | * |
| MCC950 | 0,0007 | *** |
| Time x MCC950 | 0,0559 | ns |

**G**

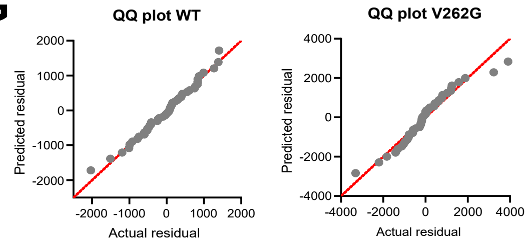


**H**

| | WT | | V262G | |
|---|---|---|---|---|
| Time (h) | p-value | p-value summary | p-value | p-value summary |
| 24 | <0,0001 | **** | <0,0001 | **** |
| 30 | <0,0001 | **** | 0,1715 | ns |
| 36 | 0,003 | ** | 0,526 | ns |
| 42 | <0,0001 | **** | 0,0165 | * |

**I**

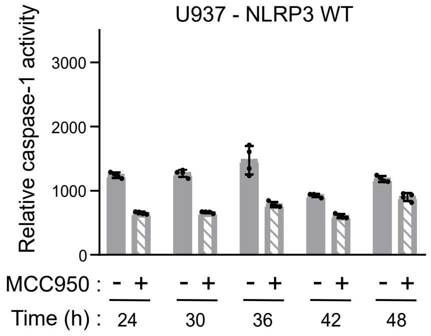


| 2-way ANOVA | p-value | p-value summary |
|---|---|---|
| Time | 0,0009 | *** |
| MCC950 | <0,0001 | **** |
| Time x MCC950 | <0,0001 | **** |

**J**

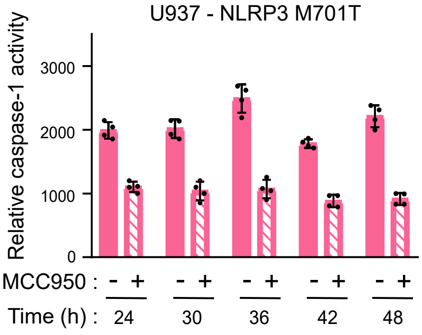


| 2-way ANOVA | p-value | p-value summary |
|---|---|---|
| Time | <0,0001 | **** |
| MCC950 | <0,0001 | **** |
| Time x MCC950 | <0,0001 | **** |

**K**

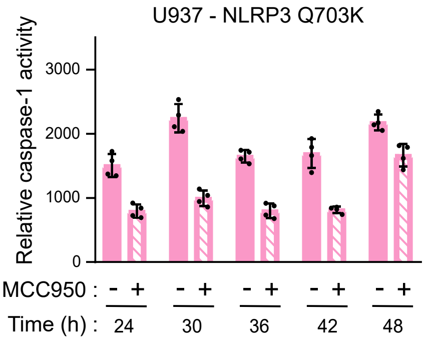


| 2-way ANOVA | p-value | p-value summary |
|---|---|---|
| Time | <0,0001 | **** |
| MCC950 | <0,0001 | **** |
| Time x MCC950 | <0,0001 | **** |

**L**

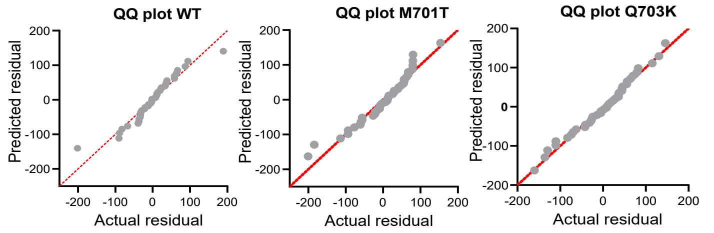


**M**

| 3-way ANOVA | M701T | | Q703K | |
|---|---|---|---|---|
| Source of variation | p-value | p-value summary | p-value | p-value summary |
| Time | 0,0002 | *** | <0,0001 | **** |
| Genotype | <0,0001 | **** | 0,3763 | ns |
| MCC950 treatment | <0,0001 | **** | <0,0001 | **** |
| Time × Genotype | <0,0001 | **** | <0,0001 | **** |
| Time x MCC950 treatment | <0,0001 | **** | 0,0001 | *** |
| Genotype × MCC950 treatment | <0,0001 | **** | <0,0001 | **** |
| Time x Genotype × MCC950 treatment | 0,0003 | *** | 0,0002 | *** |

**N**

| | WT | | M701T | | Q703K | |
|---|---|---|---|---|---|---|
| Time (h) | p-value | p-value summary | p-value | p-value summary | p-value | p-value summary |
| 24 | <0,0001 | **** | <0,0001 | **** | <0,0001 | **** |
| 30 | 0,0005 | *** | <0,0001 | **** | <0,0001 | **** |
| 36 | 0,0357 | * | <0,0001 | **** | <0,0001 | **** |
| 42 | <0,0001 | **** | <0,0001 | **** | <0,0001 | **** |
| 48 | 0,0025 | ** | <0,0001 | **** | 0,0002 | *** |
